# Perceived ambiguity of social interactions increases coupling between frontal and temporal nodes of the social brain

**DOI:** 10.1101/2020.03.11.987792

**Authors:** Matthew Ainsworth, Jérôme Sallet, Olivier Joly, Diana Kyriazis, Nikolaus Kriegeskorte, John Duncan, Urs Schüffelgen, Matthew FS Rushworth, Andrew H Bell

## Abstract

Social behaviour is coordinated by a network of brain regions, including those involved in the perception of social stimuli and those involved in complex functions like inferring perceptual and mental states and controlling social interactions. The properties and function of many of these regions in isolation is relatively well understood but little is known about how these regions interact whilst processing dynamic social interactions. To investigate whether social network connectivity is modulated by social context we collected fMRI data from monkeys viewing “affiliative”, “aggressive”, or “ambiguous” social interactions. We show activation relating to the perception of social interactions along both banks of the superior temporal sulcus, parietal, medial and lateral PFC and caudate nucleus. Within this network we demonstrate that fronto-temporal connectivity are significantly modulated by social context. Crucially, we link the observation of specific behaviours to changes in connectivity within our network. Viewing aggressive or affiliative behaviour was associated with a limited increase in temporo-temporal and premotor-temporal connectivity respectively. By contrast, viewing ambiguous interactions was associated with a pronounced increase in cingulate-cingulate, temporo-temporal, and cingulate-temporal connectivity. We hypothesise that this widespread network synchronisation occurs when cingulate and temporal areas coordinate their activity when more difficult social inferences are made.

---

Most primates live in complex social environments with large, hierarchically organized groups. Maintaining relationships within these groups impacts on individuals’ fitness (Schulke, Bhagavatula, Vigilant, & Ostner, 2010) and requires the ability to both understand the intentions and predict the actions of other individuals within the group. Recent research has identified specific brain regions that appear specialised for different aspects of social cognition and reflect the complexity of a species’ social environment (Dunbar & Shultz, 2007; Kudo & Dunbar, 2001).

These regions range in function and complexity. There are several regions in the frontal and temporal cortices that are involved in the perception of social cues such as facial expressions, body postures, and vocalisations (Bell, Hadj-Bouziane, Frihauf, Tootell, & Ungerleider, 2009; Bell et al., 2011; Diehl & Romanski, 2014; Downing, Jiang, Shuman, & Kanwisher, 2001; Downing, Peelen, Wiggett, & Tew, 2006; Hadj-Bouziane, Bell, Knusten, Ungerleider, & Tootell, 2008; Kanwisher, McDermott, & Chun, 1997; McCarthy, Puce, Gore, & Allison, 1997; Peelen, Wiggett, & Downing, 2006; Pinsk et al., 2009; Popivanov, Jastorff, Vanduffel, & Vogels, 2012; Romanski & Diehl, 2011; Scalaidhe, Wilson, & Goldman-Rakic, 1999; Sergent, Ohta, & MacDonald, 1992; Tsao, Moeller, & Freiwald, 2008; Tsao, Schweers, Moeller, & Freiwald, 2008). By contrast, there are regions concentrated in the frontal cortex and also in subcortex involved in more complex aspects of social cognition, such as the evaluation of social rewards (Aharon et al., 2001; Azzi, Sirigu, & Duhamel, 2012; Izuma, Saito, & Sadato, 2008; Rudebeck, Buckley, Walton, & Rushworth, 2006; Sescousse, Li, & Dreher, 2015; Watson & Platt, 2012), monitoring the performance of and learning from conspecifics (Behrens, Hunt, & Rushworth, 2009; Behrens, Hunt, Woolrich, & Rushworth, 2008; Kyoko Yoshida, Saito, Iriki, & Isoda, 2011; K. Yoshida, Saito, Iriki, & Isoda, 2012), and encoding of intentions and mental states of others (Haroush & Williams, 2015; Saxe & Kanwisher, 2003; Saxe, Xiao, Kovacs, Perrett, & Kanwisher, 2004; Wagner, Haxby, & Heatherton, 2012; Wagner, Kelley, Haxby, & Heatherton, 2016; Wittmann, Lockwood, & Rushworth, 2018).

The connectional properties of these social brain regions have been found, based on their activity at rest to be well preserved between humans and rhesus macaques (Mars, Sallet, Neubert, & Rushworth, 2013; Sallet et al., 2013, Mantini et al 2011). However, these studies only used resting-state fMRI to investigate the neuroanatomical properties of those networks. There is a growing interest in how these regions interact to form functional networks specialised for social behaviour. Sliwa and Freiwald (2017) contrasted responses in the monkey brain to both social interactions between conspecifics and interactions between inanimate objects lacking social associations. They identified a large “social interaction network” that included regions across the frontal, parietal, and temporal cortices as well as subcortical areas (caudate and amygdala). Intriguingly, a smaller network primarily composed of frontal regions was only responsive to social interactions and was largely unresponsive to non-social conditions.

In the human, Arioli and Canessa (2019) proposed the existence of a similar “social interaction perception” network based on the overlapping characteristics of well-established networks associated with action observation (Gallese, Fadiga, Fogassi, & Rizzolatti, 1996; Rizzolatti & Sinigaglia, 2010) and mentalising (Theory of Mind) (Koster-Hale & Saxe, 2013; Molenberghs, Johnson, Henry, & Mattingley, 2016). This network includes many of the regions mentioned above, including the posterior superior temporal sulcus, temporoparietal junction, medial prefrontal cortex, and the amygdala.

Clarifying the properties of social brain networks is of great interest, as these networks have been linked to clinically-relevant disruptions to social behaviour (e.g., autism spectrum disorders, (Liao et al., 2010); schizophrenia (Ebisch et al., 2018; Jimenez, Riedel, Lee, Reavis, & Green, 2019; Viviano et al., 2018); social anxiety disorder (Rabany et al., 2017; Zhu et al., 2017)). Despite this the majority of studies describing these social networks have either focused on the activities of individual nodes or examined the state of these networks at rest. One notable exception examined connectivity between the anterior cingulate cortex and amygdala in the gamma and beta frequency bands when animals made social decisions (Dal Monte et al., 2020). However, this study was limited to connectivity between two nodes (anterior cingulate cortex or ACC and amygdala) during a social decision. It therefore remains relatively unclear how functional relationships between nodes in social brain networks interact during the perception and evaluation of others social interactions.

In the present study, we sought to address two questions concerning neural responses to social interactions in the monkey brain: 1) how activations and network interactions are affected during the online viewing of social interactions, and more importantly, 2) how these dynamics change with respect to the nature of these interactions.

We collected functional MRI (fMRI) data from monkeys whilst they freely-viewed video clips of different social interactions between non-human primates. These social interactions included situations where the context was clear (e.g., aggressive, affiliative/grooming) and situations where the nature of the interaction was ambiguous (e.g., approach). This approach allowed us to explore changes in functional connectivity between regions responsive to social stimuli in order to better understand how the social brain functions as a network.

## RESULTS

To characterise the relationships between regions of the monkey brain involved in representing social interactions, we presented videos containing conspecific and visually-similar and closely-related but non-rhesus macaque (Macaca radiata, i.e., Bonnet Macaques) actors to three rhesus macaques whilst collecting BOLD fMRI data. The videos consisted of 5-20s clips interspersed with blank periods (Figure 1A). Each clip contained monkey actors engaged in natural behaviour with the number, identity and behaviour of the actors as well as the scene location changing randomly between clips (see METHODS AND MATERIALS for additional detail). All three monkeys were rewarded for maintaining their gaze within the borders of the video but were allowed free eye movement within this limit. On average the three monkeys maintained this level of fixation for 90±3% (M1), 89±7% (M2) and 62±8% (M3) of presented video content for each session.

**Figure 1.**
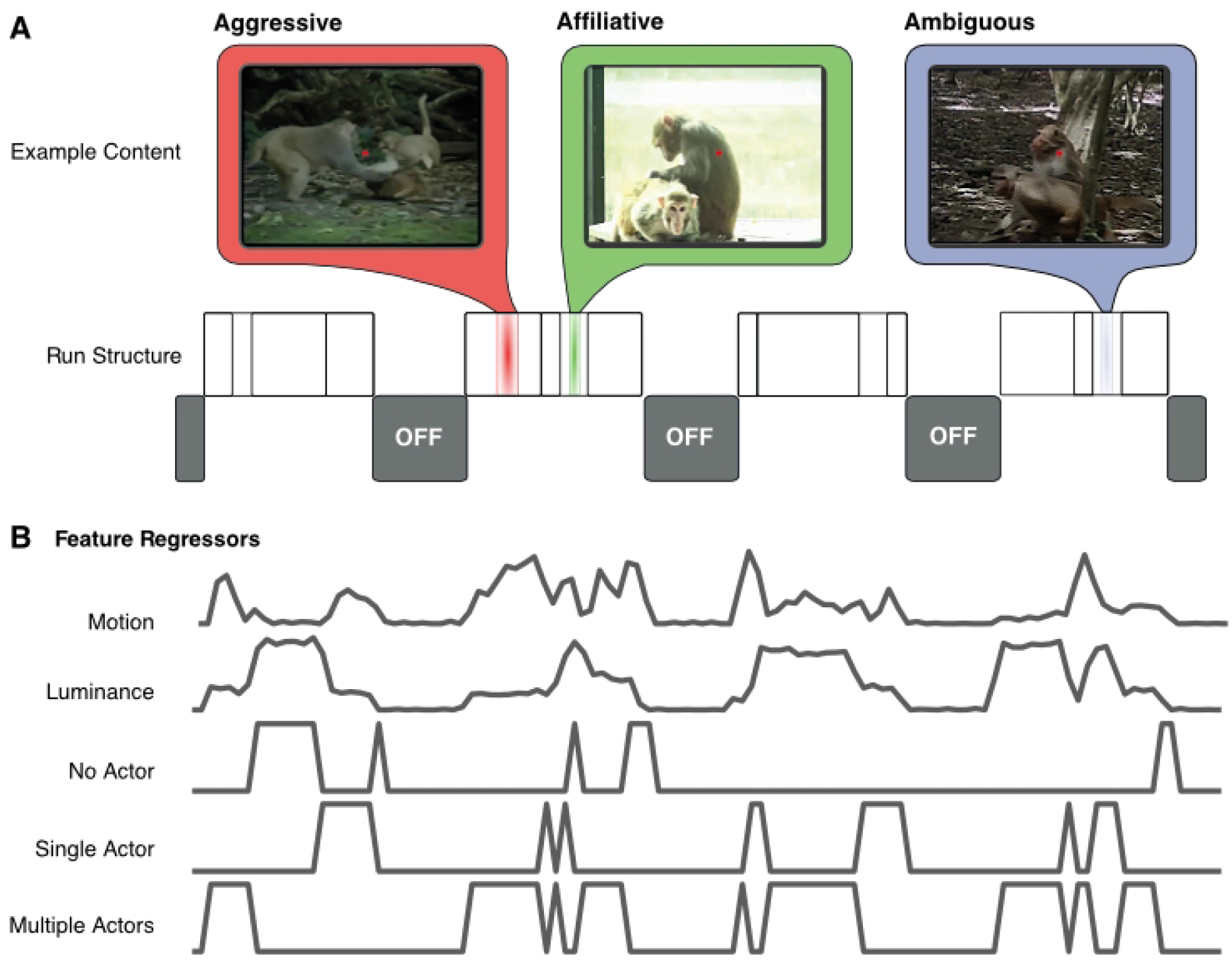
Video structure and feature regressors. **A.** Individual runs consisted of 4 video sequences interleaved with periods of blank. Each video sequence consisted of 5-20 s long clips of macaques engaged in social and non-social behaviours. Social behaviours were classified as either aggressive interactions (red), affiliative interactions (green) or ambiguous behaviour (blue). In each video sequence, periods with several immediately-abutted video clips alternated with 20 s blank periods (labelled “OFF”). **B.** Example regressors used in a GLM analysis to localise visual and social activity in the brain. Regressors were calculated from the video content and included visual features (video clips ON/OFF, luminance and motion) and social features (number of macaques present in each scene). Note regressors shown prior to convolution with the haemodynamic response function.

### Regions in the Primate Brain responsive to Social Behaviours

A univariate general linear model (GLM) analysis was conducted to identify regions in the brain that selectively respond to social stimuli (see METHODS AND MATERIALS). This model included regressors based on low-level visual features (ON/OFF, luminance and motion) as well as regressors scoring the number of actors on the screen (Figure 1B).

When monkeys viewed scenes with only single actors visible, we observed strong bilateral activation in the temporal cortex (Figure 2A; single actor>no actor, z-stat>1.9, cluster corrected, p<0.05). This activation followed the fundus of the superior temporal sulcus (STS). Within this sulcus, three semi-distinct clusters were arranged along the anterior-posterior direction and extended onto both the superior and inferior banks of the sulcus. No activation was evident outside the temporal cortex.

**Figure 2.**
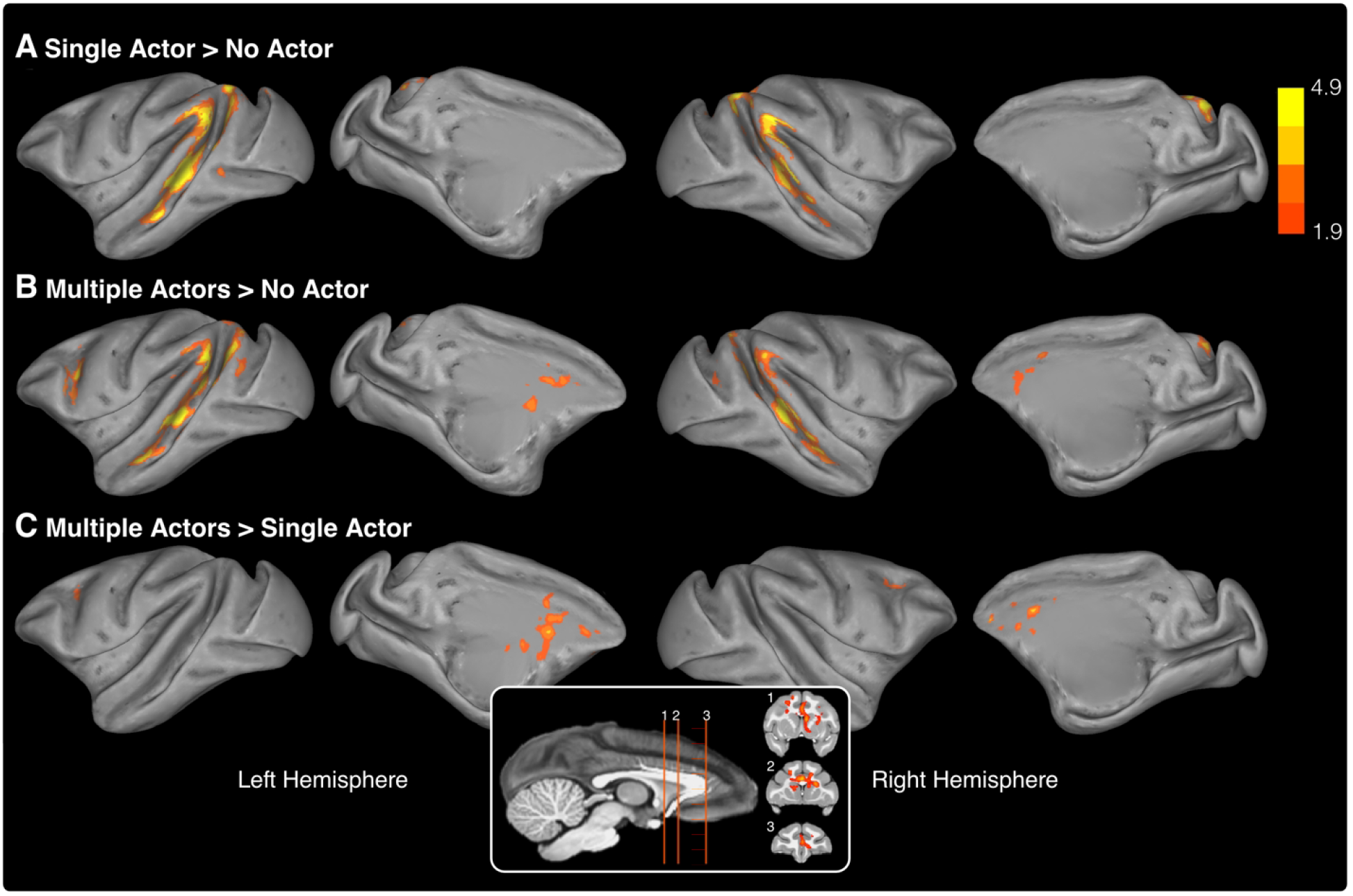
Cortical activation on viewing single actors and multiple actors engaged in natural behaviour. **A-C.** Inflated brains showing significant clusters from three contrasts derived from the number of actors visible in the videos. All data presented are from the third level, GLM analysis with combining activation from all three animals. The contrasts include; scenes with a single actor vs. scenes with no actors visible **(A)**, scenes containing multiple actors vs. scenes with no actors visible **(B)**, and scenes containing multiple actors vs. scenes containing single actors, regardless of the behaviour of the visible actors **(C)**. Medial frontal lobe activation from the multiple vs. single actor contrast is shown inset overlaid on coronal anatomical slices. All data shown survived a cluster correction at z-stat>1.9 and p<0.05.

By contrast, when monkeys viewed scenes featuring more than one actor regardless of their behaviour, strong activation was observed in both the frontal and temporal cortices (multiple actors > no actors; Figure 2B). Within the temporal cortex, activation was again bilateral and closely matched the STS clusters observed when monkeys viewed scenes containing single actors. Activation within the frontal lobe was less extensive and limited to two discrete clusters. The larger cluster extended bilaterally along the cingulate gyrus while the smaller cluster was located around the spur of the arcuate sulcus in the left premotor cortex.

Directly contrasting the responses for single vs. multiple actors (irrespective of behaviour) revealed strong activation within the medial frontal lobe (multiple actors>single actor; Figure 2C). This contrast showed bilateral activation within the cingulate gyrus and extended in the left hemisphere into the caudate nucleus (z-stat>1.9, cluster-corrected p<0.05). The three regressors of no-interest that accounted for the low-level visual features (video onset/offset, motion and luminance) predictably elicited strong activation along both banks of the STS and to a lesser extent in the tertiary visual areas (Figure S1, note differing scales).

### Dynamic connectivity within a Social Network

The above data show that activation of prefrontal areas occurred only when monkeys viewed scenes containing multiple monkeys. We therefore examined the functional interactions in the form of dynamic connectivity between frontal, temporal and subcortical regions corresponding to instances where monkeys viewed the different social behaviours. We did so using a progressive four-stage analysis approach and present the results of each stage in order to clearly illustrate how the final analysis was achieved.

Briefly, we first assessed the suitability of this approach in identifying how social behaviours affect global connectivity within the network by averaging connectivity measures across all ROIs and social behaviours (Stage 1). Next, we examined changes to individual pairwise-connections in response to non-social stimuli (Stage 2) followed by changes in response to any of the social behaviours (Stage 3). Finally, we examined what specific behaviours elicited the most notable changes to pairwise connections within the network (Stage 4).

#### (1) Global changes in response to social behaviours

We first defined a “putative social network” consisting of 8 bilateral ROIs (total 16) from clusters identified in the previous analysis. These ROIs were centred on the maximally-responsive voxels identified within the contrasts of interest above and included locations in the temporal cortex (three ROIs located along the STS), the parietal cortex (area 7a), the cingulate gyrus (incl. area 24a/b), premotor cortex, and the caudate nucleus (Figure 3A).

**Figure 3.**
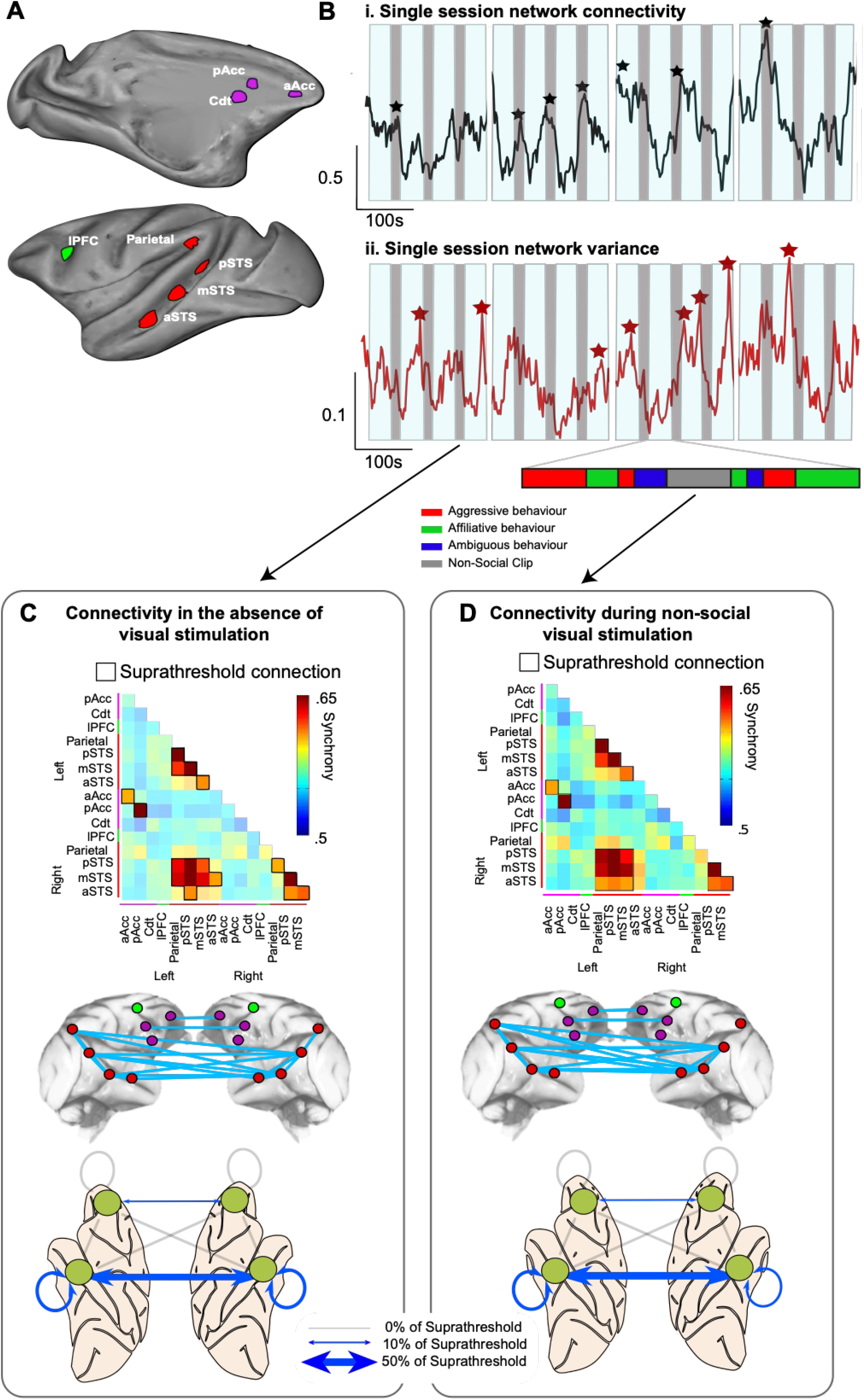
The structure and dynamic connectivity of the putative social network. **A.** Lateral and medial views of a single inflated hemisphere showing the 8 ROIs selected from the z-stat maps in Figure 2 as constituting the core of a social network. Note the colour scheme for mPFC (purple), lPFC (green) and temporal/parietal ROIs (red) **B.** Single session example**s** of the global dynamics of the network. The average dynamic connectivity between ROIs in the network, calculated using a time-windowed phase synchrony measure (*top*, black trace, calibration 100s and 0.5 AU) and the average variance in connectivity within the network (*lower*, red trace, calibration 100s and 0.1 AU). Both examples were averaged over the 880s of unique video content presented in a single session. The ON/OFF structure of the video is shown behind each trace (movie ON/OFF denoted by light blue/grey bars respectively). Interruptions in the bars denote the stop/start of each of the four individual runs. **C-D.** Detailed analysis of the structure of the putative social network in the absence of visual stimulation **(C)** and during non-social visual stimulation **(D)**. Connectivity matrices (*top*) show the strength of all possible connections between ROIs during both these conditions. Suprathreshold connections (the strongest 15% of connections, outlined in black) were selected from both matrices and the anatomical properties of the connections visualised with two network schematics. In the first schematic suprathreshold connections (shown in light blue) are displayed linked tolinking the relevant ROIs (coloured according to the above scheme) of the core network (*middle*). In addition, suprathreshold connections are summarised in a simplified representation linking the left and right frontal and temporal lobes. The thickness of the connection between these lobes corresponds to the proportion of the total suprathrehsold connections which are present between the lobes (*lower*).

Dynamic connectivity (see METHODS AND MATERIALS) was calculated between all possible pairings of ROIs in this network over an entire session. After preprocessing and concatenating the individual BOLD time-series for each run, time-series were averaged over videos repeated within a session and the resulting changes in connectivity were examined relative to both the visual and social features of the 880-s video sequence (see Figure S2 and METHODS AND MATERIALS for more details).

We first focused on two measures of connectivity within this network: 1) changes in the average connectivity, calculated across all pairwise connections within the putative social network; and 2) changes in the variance in connectivity, again using the same approach.

Both of these measures varied considerably over the time-course of the videos with sharp, transient increases in both average connectivity and variance (Figure 3Bi). Peaks in the average network connectivity were generally time-locked with periods during which the monkeys were required to maintain fixation in the absence of visual stimulation (black markers). By contrast, peaks in network variance occurred predominantly during periods of visual activation (red markers, Figure 3Bi & ii).

This latter point raises the possibility that either the visual and/or social content of the video clips was associated with changes in connectivity across a smaller number of specific connections, rather than a more uniform network-wide change in connectivity (which would have presumably increased the average connectivity, but not the variance).

#### (2) Specific changes to pairwise connections in response to non-social stimuli

We therefore performed a similar analysis, this time focussing on specific pairwise connections within the network. This analysis revealed that certain periods were marked by selective increases in specific network connections (Figure 3C & D). During blank periods (no video clips), the network was dominated by temporo-temporal connections (Figure 3C). Suprathreshold connections (which we have here defined as the strongest 15% of all pairwise connections) were primarily interhemispheric, temporo-temporal connections (44% of suprathreshold connections), followed by within-hemisphere connections in both right and left temporal cortices (22% and 22%, respectively), as illustrated in Figure 3C, right.

By contrast, connectivity within the frontal cortex was less evident (accounting for 10% of suprathreshold connections). These connections were largely inter-hemispheric, linking left and right cingulate cortex (Figure 3C). No intra-hemispheric fronto-frontal connections or fronto-temporal connections were defined as suprathreshold.

During periods with non-social video clips (visual scenes lacking any monkey actors), network connectivity was again dominated by temporo-temporal connections (Figure 3D). These primarily included interhemispheric temporo-temporal connections (50% of suprathreshold connections) followed again by within-hemisphere connections in both right and left temporal cortices (17% and 22%, respectively). Again, connections involving frontal regions were less affected (only 11% of suprathreshold connections involved areas of the frontal lobe) with the only suprathreshold connections being those linking left and right cingulate cortex, as illustrated in Figure 3D, right. No intra-hemispheric fronto-frontal connections or fronto-temporal connections were suprathreshold.

#### (3) Prefrontal-temporal connectivity is modulated by social information

To assess changes in network connectivity associated with viewing specific behaviours, we used a repeated measure ANOVA consisting of one between-subject factor: monkey (three levels, monkeys M1-M3), and one within-subject factor of interest: social interactions (three levels aggressive/affiliative/ambiguous, see METHODS AND MATERIALS for details). For this analysis, we applied a statistical threshold (z>1.66) to the matrix of z-stats for social interactions to identity connections of interest.

This approach revealed that social behaviour was correlated with modulation of predominantly fronto-temporal connections (Figure 4). Intrahemispheric fronto-temporal connections accounted for 34% of total suprathreshold connections (17% of connections for the left and right hemispheres). Interhemispheric fronto-temporal connections accounted for an additional 34% of total suprathreshold connections. By contrast, fewer suprathreshold connections were located solely within either the frontal or temporal lobes. Suprathreshold fronto-frontal connections (both linking left and right frontal lobes and within the right frontal lobe) accounted for 16% of total suprathreshold connections. Suprathreshold temporo-temporal connections were also sparse and accounted for a further 16% of total suprathreshold connections. These included intra-hemispheric connections within the left and right temporal lobes and interhemispheric connections (11, 0 and 5% of total suprathreshold connections respectively).

**Figure 4.**
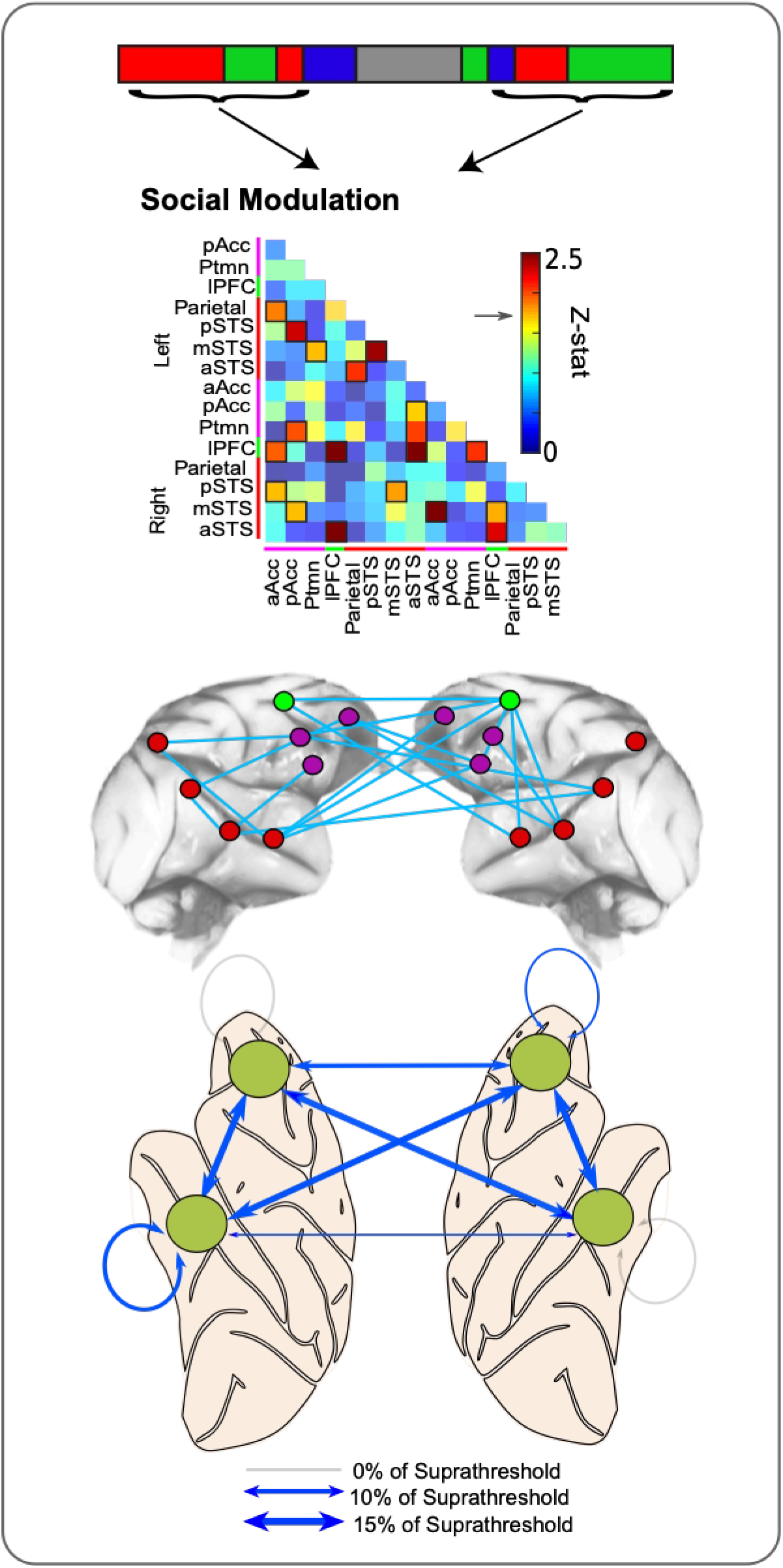
Social modulation of network connectivity. Results of a repeated measure ANOVA assessing the degree to which network connectivity was modulated by social interaction (within-subject factor with three levels aggressive/affiliative/ambiguous, See METHODS AND MATERIALS for details). The z-stat obtained from this analysis for each connection is displayed in a summary matrix (*top*). Connections with the strongest social modulation were selected with a threshold of z>1.66 (equivalent to the strongest 15% of connections, suprathreshold connections outlined in black). Suprathreshold connections are graphically represented in blue between ROIs in the network (*middle*). Simplified graphical representation of connections between the left and right frontal and temporal lobes. The thickness of the connecting line denotes the proportion of suprathreshold connections displaying social modulation of connectivity (*lower*).

To further examine which specific ROIs were linked by suprathreshold, socially-modulated connections, we calculated two metrics: degree centrality and eigenvector centrality. These were used to quantify how “central” each ROI is to the network (Figure 5). Both measures use an ROI’s connectivity to indicate its importance to a network; an ROI’s degree centrality simply reflects the sum of its connections, while an ROI’s eigenvector centrality gives greater weights to nodes connected to other well connected nodes (see MATERIALS AND METHODS for more detail).

**Figure 5.**
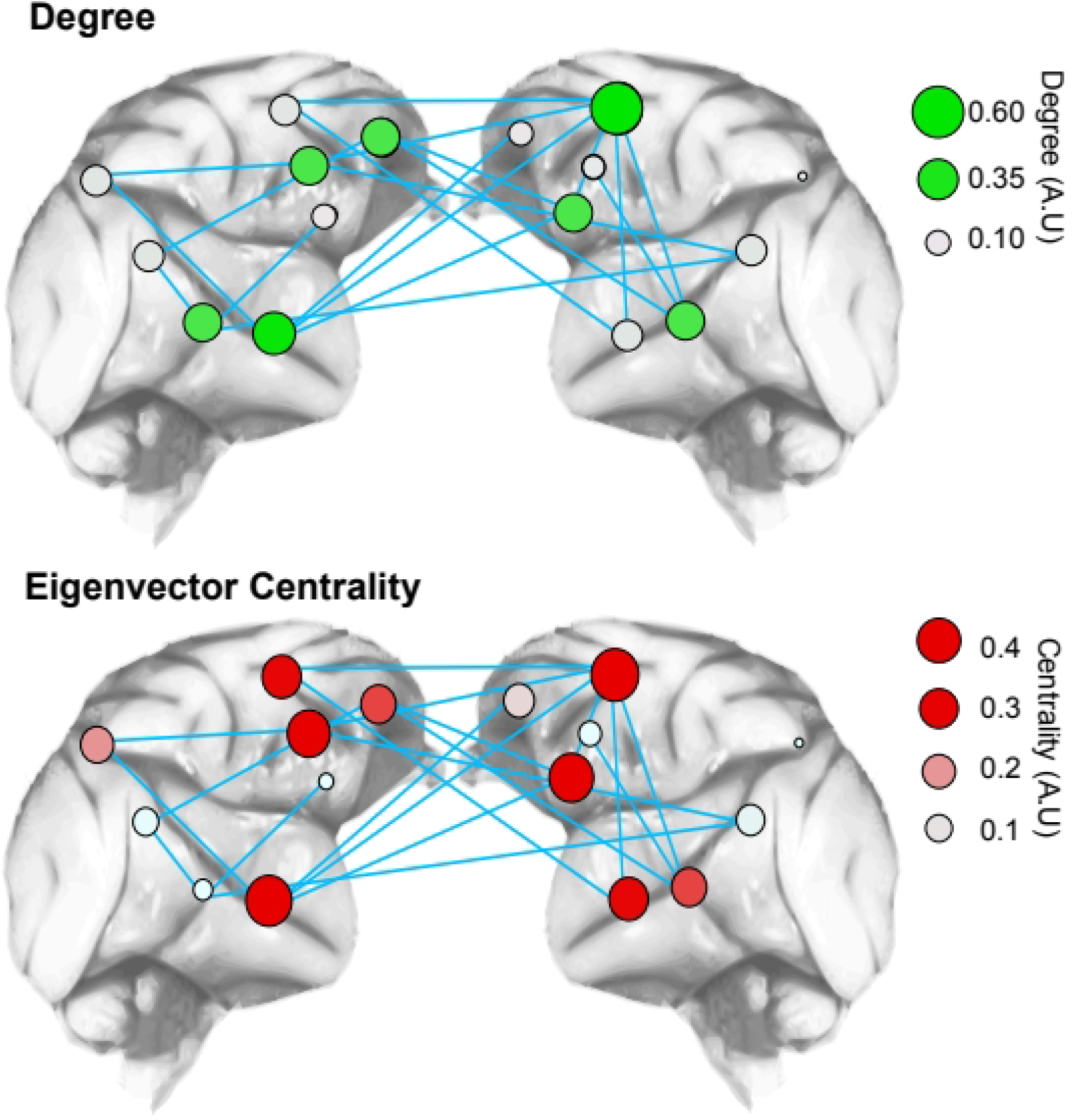
Network degree and eigenvector centrality of suprathreshold socially modulated connections. Graphical representation of degree (*top*) and eigenvector centrality (*bottom*) of each ROI calculated from suprathreshold socially modulated connections. ROI’s with greater degree (green nodes) and centrality (red nodes) indicated as larger and stronger coloured nodes within the network.

This analysis revealed a clear distinction within the temporal lobe. In contrast to posterior temporal ROIs, those ROIs more anterior along the superior temporal sulcus (aSTS & mSTS) exhibited both a greater degree (therefore more likely to be linked with suprathreshold social modulated connections) and greater centrality (more likely to be connected to other ROIs with a high number of socially modulated connections). No such distinction was evident within the frontal lobe. Although ROIs within the cingulate gyrus (notably those in the left hemisphere) exhibited both high degree & centrality scores the same was true for the premotor cortex ROI (particularly the right premotor cortex ROI).

#### (4) Ambiguous scenes elicit increased functional connectivity in fronto-temporal connections

The above data demonstrate that connectivity, both within the frontal lobe and between the frontal and temporal lobes is modulated by the nature of social behaviour viewed by a monkey. This analysis does not reveal which specific behavioural types contained in the video sequences (aggressive, affiliative, or ambiguous behaviour) were associated with the observed changes in connectivity – a vital question.

Previous work has shown how it is possible to link specific changes in network correlations with specific scenes in a video (Russ & Leopold., 2015, Hasson et al., 2004). We used a similar approach to examine how connectivity changes in response to the three types of behaviour (Figure 6A). We aligned the average connectivity between ROIs in the frontal and temporal lobes to the onset of the video clips and normalised these values to the prior baseline to show relative changes in connectivity (see MATERIALS AND METHODS & Figure S2, Stage 3). As the previous analysis indicated the existence of socially modulated connections to both cingulate gyrus and premotor ROIs, we explicitly averaged connectivity between both these sets of ROIs and the temporal lobe (Figure 6B upper and low panels respectively).

**Figure 6.**
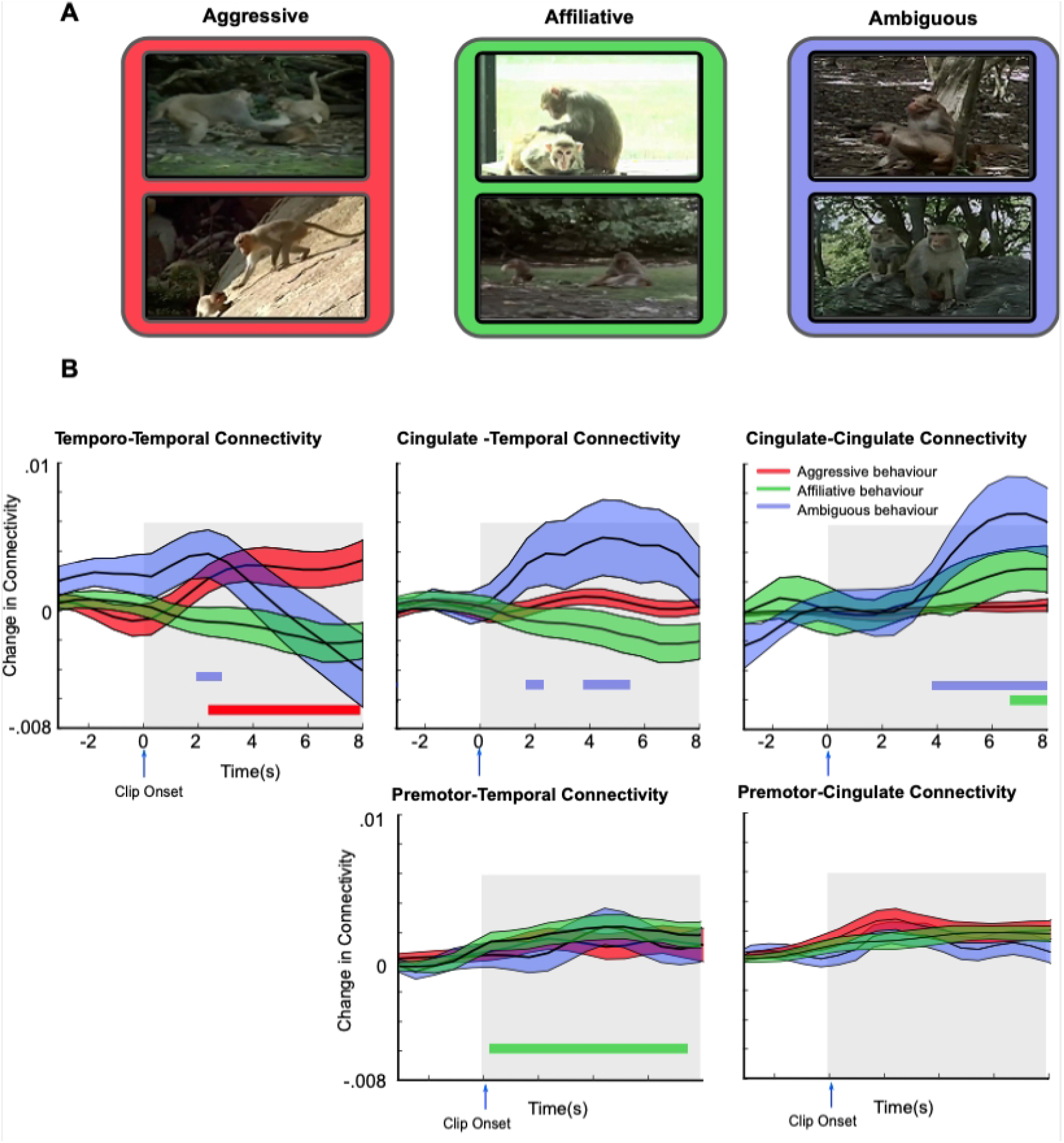
Viewing ambiguous behavioural interactions drives increased connectivity between cingulate gyrus and temporal lobe. The average time-course of connectivity between the cingulate gyrus and premotor cortex and temporal lobe ROIs aligned to the onset of clips in which the behavioural interactions were classified as either aggressive (red), affiliative (green) or ambiguous (blue). **A.** Example frames of the three behaviours contained in the clips. **B.** The clip-onset triggered connectivity was calculated from temporo-temporal (*upper left panel*), cingulate-temporal (*upper middle panel*), cingulate-cingulate (*upper right panel*), premotor-temporal (*lower middle panel*), and premotor-cingulate (*lower right panel*) connections for each of the three behaviours viewed. Consistent with previous analyses only the strongest 15% of connections were considered. Coloured bars denote timepoints with significantly increased connectivity at p<0.05. Significance was determined by one-tailed t-test against surrogate data generated with the same mean and variance as the pre-clip baseline. To correct for multiple comparisons cluster correction was applied across timepoints at p<0.05.

This analysis, shown in Figure 6, revealed differences in connectivity within our network depending on the specific behaviour viewed. Firstly, increased connectivity between the cingulate gyrus and temporal lobe was predominantly driven by video clips where the behaviour was classified as ambiguous rather than clips where the behaviour was clearly affiliative or aggressive. Specifically, viewing clips containing ambiguous behaviour was associated with significant increases in temporo-temporal, cingulate-temporal and cingulate-cingulate connectivity (Figure 6B, blue traces). By contrast, viewing clips containing aggressive behaviour was associated with significant increases in temporo-temporal connectivity only with no significant changes in cingulate-temporal or cingulate-cingulate connectivity (Figure 6B, red traces). Finally, viewing clips showing clear affiliative behaviour between monkeys was not associated with any changes in temporo-temporal or cingulate-temporal connectivity but was associated with a small significant increase in both cingulate-cingulate connectivity (Figure 6B, green traces) and premotor-temporal lobe connectivity.

## DISCUSSION

We have examined how regions of the monkey brain responsive to social stimuli coordinate their activities in response to dynamic and complex social interactions. Monkeys were shown short video clips involving one or more monkey actor and which broadly fell into one of three categories: affiliative, aggressive, and a third category where the nature of the interaction was ambiguous to the viewer (e.g., clips of one actor approaching another, or two actors approaching one another).

Viewing clips of social interactions activated several brain regions, concentrated within the frontal and temporal lobes (Figure 2). Critically, although temporal lobe regions were activated regardless of the number of actors, frontal areas were only recruited whilst monkeys viewed clips with more than one actor. Connectivity across this social brain network, and in particular between/within the frontal and temporal regions varied according to the behaviour viewed by the monkeys. Viewing affiliative behaviour was associated with a small significant increase in connectivity between premotor and temporal lobe ROIs, but had no impact on cingulate-temporal lobe connectivity. By contrast, significant increases in connectivity within the cingulate gyrus and between cingulate gyrus and temporal ROIs were observed when monkeys viewed ambiguous behaviour (Figure 6). We propose that the increase in cingulate-cingulate and cingulate-temporal lobe connectivity associated with viewing ambiguous social interactions may reflect the increased neural processing necessary to make accurate predictions about upcoming behaviours that are unnecessary or reduced when subjects view more predictable interactions.

Here, we first present how our selection of nodes in our social network relates to pre-existing literature. Before, discussing why certain social contexts may lead to increased functional connectivity between nodes in this network.

### Composition of a social network

Recent advances in the field of social cognition, including in the non-human primate brain have led to a better level of understanding of the individual roles for these regions (Platt, Seyfarth, & Cheney, 2016). For our purposes, we selected a subset of brain regions on the basis of which areas were reliably activated by our stimuli using our imaging protocols. We grouped these regions into a “putative social network”.

The vast majority if not all of the regions in our putative social network are well-established as being involved in social cognition. To begin, our social network included three bilateral ROIs along the STS, presumably corresponding to the canonical face-selective regions (Bell et al., 2009; Pinsk et al., 2009; Tsao, Moeller, et al., 2008). Although we did not independently assess the boundaries for body-part selective regions, they are typically located immediately adjacent to face-selective regions (Bell et al., 2009; Pinsk et al., 2009), and so we can assume our STS-regions incorporate body-part selectivity as well.

The contribution of these regions to our social perception task may seem self-evident. And yet, more recent structural studies have shown the STS to be sensitive to the composition and hierarchy of the monkey actors, not just their identity (Noonan et al., 2014; Sallet et al., 2011). It is possible that in addition to participating in the “simple” visual processing and reconstruction of the images (e.g., extracting the visually-derived semantic codes as proposed by (Bruce & Young, 1986), these regions are playing a more complex role in evaluating non-visual identity-derived semantic features of the actors, such as their degree of dominance.

The two ROIs located in the cingulate cortex also have a role in social cognition in both humans and non-human primates. In particular, the cingulate gyrus has been shown to be central to social valuation (Apps & Ramnani, 2014; S. W. Chang, Gariepy, & Platt, 2013; Rudebeck et al., 2006). Adjacent areas of cingulate cortex and other medial frontal regions have also been implicated in tracking the behaviour and intentions of other agents (Fatfouta, Meshi, Merkl, & Heekeren, 2018; Haroush & Williams, 2015; Hill, Boorman, & Fried, 2016; Lockwood & Wittmann, 2018; Wittmann et al., 2016; Kyoko Yoshida et al., 2011; K. Yoshida et al., 2012).

In addition, our network included a premotor ROI that was located near the spur of the arcuate sulcus. This is notable, as premotor cortex is increasingly being implicated in social cognition. For example, detailed single neuron recordings have revealed sub-populations of mirror neurons in premotor area F5 that respond to actions essential for judging social hierarchy and status including gaze direction (Coudé et al 2016) and lip smacking (Ferrari et al 2003). Furthermore, Sliwa & Freiwald recently demonstrated significant overlap between two contrasts: one mapping social interactions and the other localisation of the mirror neuron system (Sliwa et al 2017, Nelissen et al 2011), arguing that this overlap indicated a role for mirror neurons in processing social intentions of an interaction as well as simple motor understanding of the interaction.

Finally, we observed activation in the caudate nucleus. While we cannot speculate what specific role caudate may serve in interpreting social behaviour, the area’s contribution to social cognition in general is becoming more clear. It has recently been associated with the default mode network (Alves et al., 2019), which itself has been linked to the social brain (Mars et al., 2012). In humans, responses to monetary and social rewards have been associated with striatal activity (Izuma et al., 2008). In monkeys, feedback for self vs. others is discriminated by striatal neurons (Baez-Mendoza, Harris, & Schultz, 2013; Baez-Mendoza, van Coeverden, & Schultz, 2016). Closer to our protocol, Sliwa and Freiwald (Sliwa & Freiwald, 2017) found that caudate was activated during viewing of social scenes; and Noonan and colleagues (Noonan et al., 2014) found that the volume of grey matter in the caudate covaried with social status in macaques.

One notable absence in our social network was the amygdala – we did not observe statistically significant activation in the amygdala for any of our social videos, despite its well-established role in face processing and social behaviour. There are at least two possible explanations. The first is a technical limitation, which is that the amygdala is difficult to image in monkeys, particularly larger male monkeys who have extensive musculature on either side of their skulls. This additional muscle mass increases the distance between the receiver coil elements and the target structure, thus inhibiting signal detection. However, we note that previous studies which have used either the same experimental setup (Chau et al., 2015) or similar study design (Sliwa & Freiwald, 2017) have found amygdala activity. Therefore, a second explanation is that while the amygdala may have indeed been active during our task, it was equally active across all conditions and thus no significant differences were observed. This would fit with more modern studies of amygdala function that highlight its role in generalised linking of social and non-social stimuli to outcomes rather than being, for example, a “fear module” (i.e., selectively activated by specific social behaviours, contexts, or stimuli).

### Role of prefrontal recruitment in interpreting social interactions

The most notable observation in our study was the increased functional connectivity of the cingulate cortex while monkeys observed ambiguous but not clearly affiliative or aggressive social interactions between multiple actors. In the latter two cases, the monkeys viewing the video clips would not be required to make any judgments or predictions about the nature of the interaction – as it was clearly depicted in the video. On the other hand, even in a passive viewing case, it is possible that the monkeys viewing the ambiguous clips might automatically infer the ultimate outcome of the actors’ behaviour.

In this scenario one interpretation is that inferring the consequences or outcomes of ambiguous social behaviour, even without an active task component, not only requires regions of the brain where neurons encode the social features of stimuli (eg. face patches in the inferior temporal cortex) but also areas of the medial prefrontal cortex. This hypothesis is supported by electrophysiological studies of medial prefrontal cortex in non-human primates. These studies have revealed neurons within the ACC encode a range of information essential for making social decisions including; shared reward experience (Chang, Gariepy, & Platt, 2013), the actions of other animals (Kyoko Yoshida, Saito, Iriki, & Isoda, 2011;) and a prediction of other animals future decisions (Haroush & Williams, 2015). Furthermore, recent research has revealed that synchrony between neurons of the ACC and other brains areas, in this instance the amygdala, can be modulated by social context (Dal Monte et al., 2020). Dal Monte et al revealed that coherence between neuronal activity recorded from the ACC and amygdala was increased when animals shared a reward but decreased only self rewarded.

However, it should be noted that these studies detail medial prefrontal activity whilst NHPs were required to make decisions based social cue or information, rather than observing social interaction between other animals. Further, evidence for the recruitment of prefrontal cortex during the viewing of social interactions has been observed in recent FMRI studies in humans. For example, Wagner et al. (Wagner et al., 2016) had human participants passively view movie clips while undergoing fMRI. While they consistently found frontal, temporal, and parietal activation during the movie, they noted that dorsomedial PFC was most engaged during scenes that involved social interactions between characters in the movies; raising the possibility that different regions are active during scenes where some behavioural assessment may be necessary.

Sapey-Triomphe et al. (Sapey-Triomphe et al., 2017) presented point-light images of people either interacting or not interacting to participants ranging in age from 8-41 years of age. The participants were required to state whether the images were interacting or not. Across all age groups, STS, middle temporal gyrus, anterior temporal lobe, and inferior frontal gyrus were activated during the social interactions. However, stronger activation was observed in frontal, parietal, and striatal (caudate) areas – that is, structures implicated in mentalising behaviour – in adults, who were also more successful at classifying the point-light behaviours. Finally, Gardner et al. (Gardner, Goulden, & Cross, 2015) demonstrated that increased familiarity with videos of subjects performing dance moves resulted in decreased correlations within an action-observation network.

Collectively, these results highlight two potential features of dynamic interactions in social networks. First: that interactions between frontal, parietal, and temporal cortices are not static but rather change dynamically depending on the features and nature of the social stimuli being presented. Second, making inferences or active interpretations of social interactions (rather than passive viewing of social scenes) appears to recruit additional frontal activation.

Our data parallel these conclusions by showing the greatest amount of coherence among nodes in our putative social network when monkeys were viewing ambiguous social scenes, as compared to predictable scenes. Therefore, we speculate that this additional frontal recruitment reflects the additional cognitive demands of deciphering ambiguous social scenes.

### Future Directions

This study represents an early step towards understanding the role of frontal, striatal and temporal cortex in social cognition. Critically, our task was a passive task – the monkeys were not required to make judgements about the behaviour of the actors. This limits our ability to guess what information might be passing between the frontal and temporal regions. Yet, there is an increasing number of studies examining social behaviour between more than one subject (Grabenhorst, Báez-Mendoza, Genest, Deco, & Schultz, 2019; Haroush & Williams, 2015; K. Yoshida et al., 2012), so it is becoming more feasible to conduct experiments involving two or more monkeys interacting with one another. To fully understand how brain regions involved in social cognition interact and what type of information passes between them will likely require such experiments. Future experiments will seek to add a behavioural / decision-making component to these type of experiments – for example, having the animals guess what the outcome of different ambiguous situations might be; possibly having them make predictions about upcoming interactions. This way, we may take one step closer to an understanding of the degree to which non-human primates exhibit rudimentary “theory of mind”-like abilities, and what role regions such as those discussed in this study might play in that cognitive function.

Second, fMRI is limited in its ability to reveal the nature of information being passed from one region to another. This technique can identify circuits of interest to study with a more suitable method that can clarify the nature of the millisecond-by-millisecond information being passed between nodes in a complex cortical network. For example, how are the new information requirements in situations when animals are viewing ambiguous social interactions communicated to/from frontal and temporal regions? How are the neural representations within temporal cortex of the actors being updated or modulated while the scene plays out? These are questions that are perhaps better addressed using techniques with better temporal resolution compared with MRI-based approaches. No doubt, such experiments will yield exciting new insights as to the role of frontal/temporo- and striato/caudal interactions in social cognition.

## MATERIALS AND METHODS

All procedures were conducted under licenses granted by the United Kingdom (UK) Home Office in accordance with the UK Animals (Scientific Procedures) Act of 1986, after approval from the University of Oxford local ethical review panel and the UK Home Office Animal Inspectorate. All husbandry and welfare conditions complied with the guidelines of the European Directive (2010/63/EU) for the care and use of laboratory animals.

### Animals

Three adult male monkeys (*Macaca mulatta*, M1, M2 and M3), purpose-bred within the UK, were used in the study. The monkeys were aged 7-8 years old and weighed 10-13 kg at the time of data collection. All monkeys lived in large communal rooms (but with separate housing areas) with several other macaques with whom they could visually interact. Monkeys M1 and M2 were pair-housed whereas M3 was singly-housed. All monkeys were kept on a 12-hr light/dark cycle and were given free access to water on non-testing days, and at least 14-hr access to water on testing days. Veterinary staff and animal technicians performed regular health and welfare assessments, including formalized behavioural monitoring. Prior to fixation training all monkeys were implanted with MR compatible polyether ether ketone (PEEK) head-posts (Rogue Research, Montreal, CA) and ceramic screws (Thomas Recordings Gmbh) under aseptic conditions (for further details see (Bell et al., 2009; Chau et al., 2015)).

### Experimental set-up

Stimulus presentation, reward delivery and eye calibration were controlled using PrimatePy, an implementation of PsychoPy (Peirce, 2007) modified for primate research (Baumann et al., 2015; Joly, Baumann, Balezeau, Thiele, & Griffiths, 2014). Stimuli were projected onto a screen placed 19 cm in front of the monkeys. Eye position was recorded using an MR-compatible camera (MRC Systems GmbH, Heidelberg, De) and horizontal and vertical eye position were down-sampled to 25Hz and stored for offline analysis along with reward delivery timings and TR pulse count.

### Stimuli and data collection

We used four different 220 s video sequences (total video length 880 s) presented in separate runs. For each sequence, there were between three and four runs per daily session. Each video sequence consisted of sixteen clips of either 5,10 or 20 s long, interspersed with 20 s blank sections. A fixation cue (red circle, 0.3 deg) was visible during both video clips and blank periods of the sequences (Figure 1A). Throughout the sequences, monkeys were rewarded for maintaining their gaze within the boundaries of the videos (largest dimension set to 13 deg). Clips used in the video sequences depicted either conspecifics or the Bonnet macaque (*Macaca radiata*), a closely related species within the *Macaca* group. The clips were obtained from three sources: two nature documentaries and a set of clips filmed within a local breeding centre. Clips featuring monkeys contained either 1 or 2+ actors engaging in different social behaviours. Prior to data collection all monkeys were trained using alternative video footage consisting of sporting events. During data collection M1 participated in 12 sessions (total volumes: 18,040), M2, 12 sessions (total volumes: 16,720) and M3, 11 sessions (total volumes: 15,400).

### MRI Data acquisition

Imaging data were collected using a 3T MR scanner and a four-channel phased-array receive coil in conjunction with a radial transmission coil (Windmiller Kolster Scientific, Fresno, USA). Both fMRI images and proton-density weighted reference images were collected while awake animals were head-fixed in a sphinx position in an MR-compatible chair (Rogue Research, Montreal, CA). fMRI data were acquired using a gradient-echo T2* echo planar imaging (EPI) sequence with 1.5×1.5×1.5 mm resolution, 32 ascending slices, TR = 2 s, TE = 29 ms, flip angle = 78. Proton-density weighted images using a gradient-refocused echo (GRE) sequence (TR=10 ms, TE= 2.52 ms, flip angle= 25) were acquired as reference for body motion artefact correction during pre-processing. T1-weighted MP-RAGE images (0.5×0.5×0.5 mm resolution, TR = 2,500 ms, TE = 4.01 ms, 3-5 sequences per image) were acquired from each of the three monkeys in separate scanning sessions and were collected under general anaesthesia (see (Ainsworth et al., 2018; Mitchell et al., 2016) for further details of anaesthesia protocols and T1 image acquisition).

### Data Analysis

#### fMRI pre-processing

Initial fMRI data pre-processing was carried out on a run-by-run basis using Matlab toolboxes developed to correct for common artefacts in monkey functional imaging (Offline Sense and Align EPI toolboxes, Windmiller Kolster Scientific, Fresno, USA). Data were first reconstructed offline from raw image files using SENSE reconstruction to reduce Nyquist/ghost artefacts (Kolster et al., 2009). Non-linear motion artefacts in the data were corrected on a slice-by-slice basis using a 3rd order polynomial to align all volumes within a run to an ideal EPI reference image (Kolster, Janssens, Orban, & Vanduffel, 2014).

Further pre-processing of the reconstructed and motion corrected data was carried out using functions from both AFNI (Cox, 1996) and FSL (fMRI of the Brain (FMRIB) Software Library (Jenkinson, Beckmann, Behrens, Woolrich, & Smith, 2012). Individual runs were concatenated to yield a single 4D data file for each session and the resultant data were skull stripped and signal outliers were removed (using 3dDespike from the AFNI package, (Cox, 1996)). Remaining volumes that were contaminated by excessive motion were identified based on the volume to volume variance (fsl_motion_outliers using the dvars option, (Power, Barnes, Snyder, Schlaggar, & Petersen, 2012)). For each session, individual volumes with variance greater than the session mean plus two and a half times the session standard deviation were identified as outliers and modelled in further analysis as nuisance regressors. For each monkey, the average percentage of volumes per session identified in this way were; M1 4±2%, M2 6±1% and M3 7±1%.

Data were registered to the NMT standard monkey atlas (Seidlitz et al., 2018) with a two-step registration process. First, the mean EPI image for each session was registered to the relevant monkey’s high resolution T1-weighted structural image. This was achieved by boundary-based registration of mean images, with field maps used to simultaneously correct for EPI field distortions (Greve & Fischl, 2009; Jenkinson, Bannister, Brady, & Smith, 2002; Jenkinson & Smith, 2001). Each monkey’s T1 structural image was then registered to the NMT template image within 9-degrees of freedom. For each session, the two relevant transformation matrices were combined and saved for further analysis. Segmentation of T1 structural images to generate grey, white matter and CSF masks was achieved using FAST (Zhang, Brady, & Smith, 2001) and masks for each monkey were transformed into EPI-space for use in further analysis. Finally, during initialisation of the general linear model (GLM, see below for details), fMRI data were spatially smoothed (3mm FWHM), temporally filtered (3-dB cut-off 100s), and intensity normalised.

#### Video feature regressors

Regressors coding for visual and social features of the stimuli were created on a frame by frame basis from the content of the videos (examples shown in Figure 1B). First, a binary video ON/OFF regressor was created in which ones corresponded to frames with video content and zeros for frames during blank fixation periods. Two additional regressors were created based on the overall luminance and total motion of each frame of the video. Total motion between video frames was calculated using a block matching method with video frames divided into 25×25 pixel blocks (Block Matching function, Computer Vision System Toolbox, Matlab 2014a, Mathworks). Finally, three binary regressors were created based on the number of monkeys visible on each frame (no monkeys, one monkey and two or more monkeys). Video content was manually scored on a frame-by-frame basis and assigned to the appropriate regressor. All regressors were downsampled to 0.5Hz to match the 2 s TR of the fMRI sequence. Prior to convolution with a gamma function (SD 1.5 s, mean lag 3 s), all regressors were modified such that volumes in which the monkey failed to fixate for more than 80% of that volume were set to zero.

#### Whole brain GLM analysis

We conducted an initial analysis to identify brain regions that respond selectively depending on number of actors (Figure 2). We used a multilevel, univariate GLM analysis using the FSL FEAT tool (Woolrich, Behrens, Beckmann, Jenkinson, & Smith, 2004). The first level GLM in this analysis was carried out on the processed 4D fMRI data of each session. The model at this level included regressors for low level visual features (Video ON/OFF, total motion between video frames and overall luminance) as well as the regressors of interest (number of actors, 0, 1 or multiple). Three contrasts of interest were included in the model to identify areas of the brain activated by viewing differing numbers of animals: one actor vs. no animals, multiple actors vs. no animals and multiple vs. single actors. In addition, individual regressors were included in the model for each volume identified as being contaminated by excessive motion using fsl_motion_outliers (described above). The results from the first level analyses were then combined in three, second-level mixed-effects GLM’s (FLAME 1 & 2), corresponding to one for each monkey. We then combined these into a third, final group-level analysis consisting of a further fixed-effects GLM (Woolrich et al., 2004). Significant clusters were identified from the z-stat images using a threshold of z > 1.9 and cluster-correction of p < 0.05.

#### Social network ROI definition

ROIs within our putative social network were defined based on the activation clusters for two contrasts: single>no monkeys and multiple>no monkeys. Local maximal voxels were identified within each cluster obtained from each of these contrasts. To ensure ROI size was consistent across the left and right hemispheres, all maximal voxels were projected onto a single hemisphere and the strongest eight voxels were selected. These eight voxels were then mirrored in both hemispheres and a total of 16 spherical ROIs with a diameter of 4mm were generated across the two hemispheres (8 per hemisphere). We chose to focus on only 8 ROIs per hemisphere as this gave a reasonable distribution of well-sized ROIs across frontal and temporal cortex, without including spurious clusters.

ROIs were masked according to the LV-FOA-PHT cytoarchitectonic standard atlas (Van Essen, Glasser, Dierker, & Harwell, 2012) such that each ROI was constrained to a single cortical area and there was no overlap between adjacent ROIs.

#### Calculation of dynamic connectivity

Prior to the calculation of dynamic connectivity between ROIs, the mean BOLD signal from all sessions from each ROI was filtered using a GLM incorporating two confound time-series: one generated from the CSF mask and another derived from timestamps denoting the onset of the reward pulses. The residual BOLD time-series obtained from this model was used for the subsequent connectivity analysis.

Dynamic connectivity was assessed by the pairwise calculation of relative phase synchrony between all ROIs (Rosenblum, Pikovsky, & Kurths, 1996). In contrast to correlation-based measures of connectivity, relative phase synchrony provides a measure of coherence unbiased by the amplitude of the signals. However, phase synchrony measures are sensitive to the frequency content of paired signals. Previous studies have considered both within (1:1) and across frequency (1:n) phase synchrony (Palva, Palva, & Kaila, 2005). In this study, no assumptions were made about specific frequency coupling and phase synchrony was calculated from 0.01Hz to 0.5Hz. Fourier analysis of the bold time series revealed peaks evident at 0.02Hz and 0.04Hz but all frequencies in the aforementioned range are considered in all subsequent analyses (Figure S3).

As with previous dynamic connectivity studies, phase synchrony was calculated for short overlapping windows of paired time-series. The length of sliding windows is typically limited by decreased signal-to-noise ratio and increased variability as window length decreases (Hutchison, Womelsdorf, Gati, Everling, & Menon, 2013) while others have suggested a minimum window size of 33 s is required to reveal stable modular architecture within the brain (Jones et al., 2012). Comparable window lengths have been used in previous dynamic connectivity studies of resting state activity (C. Chang, Liu, Chen, Liu, & Duyn, 2013; Hutchison et al., 2013). We therefore calculated relative phase synchrony between the instantaneous phase of each pair of signals over a 32s time window. To ensure the subsequent phase synchrony was calculated with sufficient temporal resolution to reveal changes linked to events within the videos, each window was offset by 2s and overlapping the adjacent window by 30s (Figure S2, Stage 1).

All synchrony values for each session were arcsine transformed to account for any values at the extremes. For each session, the normalised synchrony values were averaged across repeated viewings of the videos to yield a time-course corresponding to the complete 14.8 min of unique video content (Figure S2, Stage 2).

#### Statistical analysis of dynamic connectivity

Before analysing dynamic connectivity within our network relative to social behaviours, we first validated the technique. Initially, we examined global connectivity within the network over the timecourse of the scanning sessions by calculating the mean connectivity and mean variance across all pairwise connections in the network. We then calculated the mean strength of each connection during periods of non-interest (blank periods in the video and non-social content) and used a threshold selecting for the strongest 15% of connections to view the structure of the network.

To assess the extent to which viewing different social interactions modulated network connectivity, we averaged the phase synchrony values for each pairwise connection between ROIs within the network on a session-by-session basis for three different network states. These states corresponded to the manually-scored time-courses for scenes containing multiple actors engaged in three different behaviours: (1) affiliative behaviour (e.g., lip smacking, grooming behaviour, etc.), (2) aggressive/dominant behaviour (e.g., piloerection, teeth baring, and/or physical confrontation), and (3) ambiguous behaviour in which the nature of interactions between the two or more actors was unclear (average connectivity matrices for each state shown in Figure S4).

A repeated measure ANOVA was then conducted for each connection using these average connectivity values with one between-subject factor: monkey (three levels, monkeys M1-M3), and one within-subject factor: social interactions (three levels, affiliative/aggressive/ambiguous). The statistics obtained from this analysis were used to create a single matrix of z-stats for each connection. From this matrix, only connections with a z-stat>1.66 (representing the strongest 15% of total connections) were considered for further analysis. To determine essential or central nodes in the network, two measures were calculated from the resulting binary matrix of social modulated connections using the Brain Connectivity Toolbox (Rubinov & Sporns, 2010). Firstly, for each ROI, the degree or number of connections to the ROI was calculated. Secondly, the importance of each ROI was assessed by calculating eigenvector centrality. Eigenvector centrality is biased toward well-connected nodes. Therefore, ROIs with high eigenvector centrality are not only well connected within a network but have a lot of connections to other well connected ROIs.

To explicitly link changes in connectivity to the specific types of behaviour, an additional analysis was conducted in which changes in connectivity were examined after the onset of clips containing each of the three behaviour types (aggressive, affiliative, or ambiguous behaviour). We aligned 11-s segments of phase synchrony time-series (3 s pre-to 8 s post clip onset) with a 2-s delay to allow for the haemodynamic response. The aligned time series were then interpolated using a cubic spline and averaged from the strongest 15% of connections between five anatomical groupings: cingulate-cingulate connections, cingulate-temporal connections and temporo-temporal connections, premotor-cingulate connection and premotor-temporal connections (including connections within and across hemispheres). To identify statistically significant increases in synchrony relative to baseline for different behaviours, these clip-triggered averages were compared to randomised surrogate data (generated around the average mean synchrony and variance of the pre-onset triggered time-course) using one-tailed t-tests. Significant p-values were cluster-corrected to p<0.05.

---

**Figure S1.**
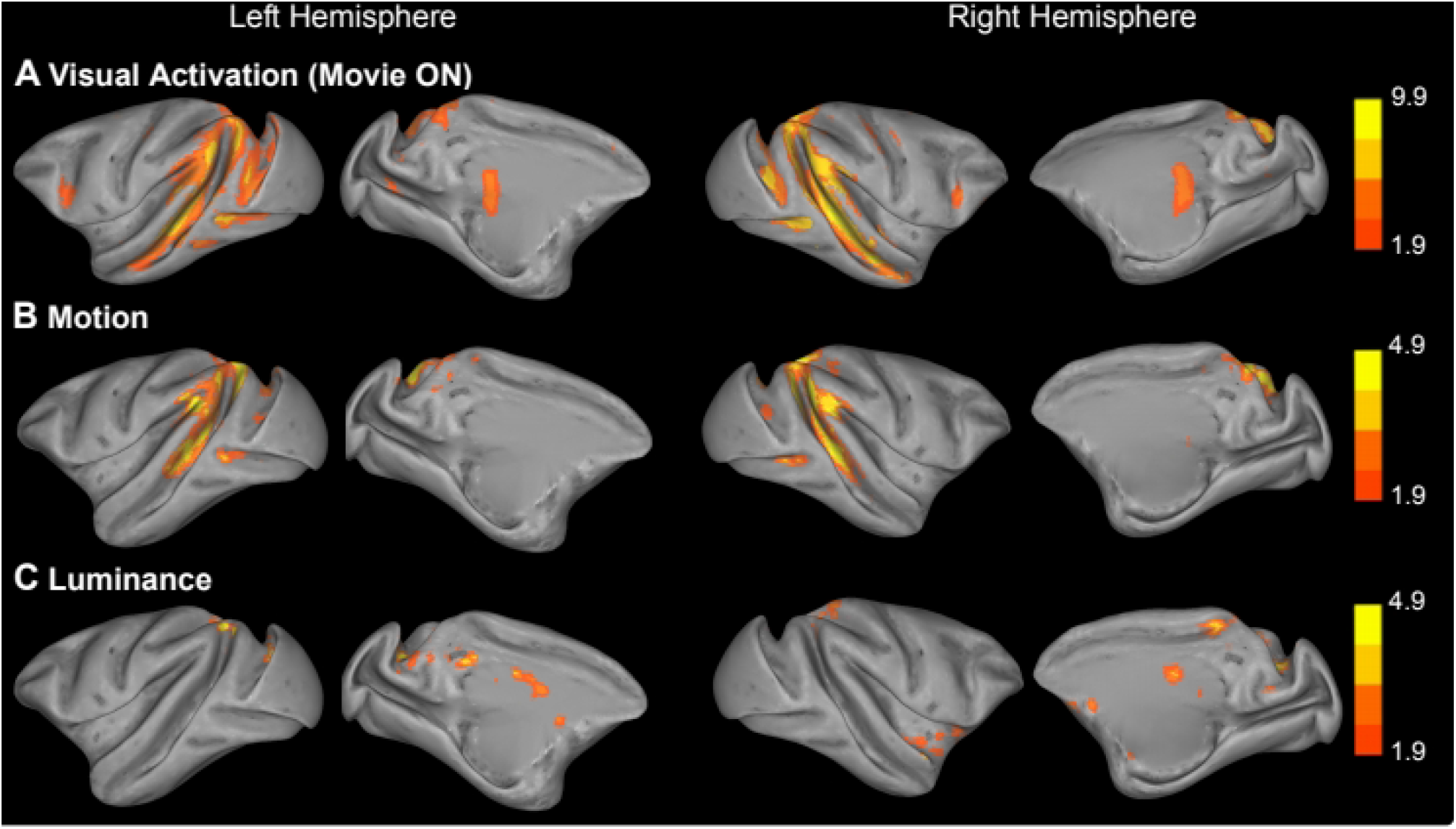
Cortical activation associated with low level visual video features. **A-C.** Inflated brains showing significant clusters from three contrasts of low level visual features calculated from the videos. All data presented are from the third level, GLM analysis with combining activation from all three animals. The contrasts include; the basic visual activation during each session (video ON/OFF, **A**), the motion within the video, calculated by a block matching algorithm examining differences between frames of the video content (See METHODS AND MATERIALS for details, **B**), and the luminance of the video scenes (**C**). Note the differences in scales as different thresholds (z-stat>6.5 z-stat>1.9 and z-stat>1.9) were applied to the data shown in A-C respectively, and all images were cluster corrected at p<0.05.

**Figure S2.**
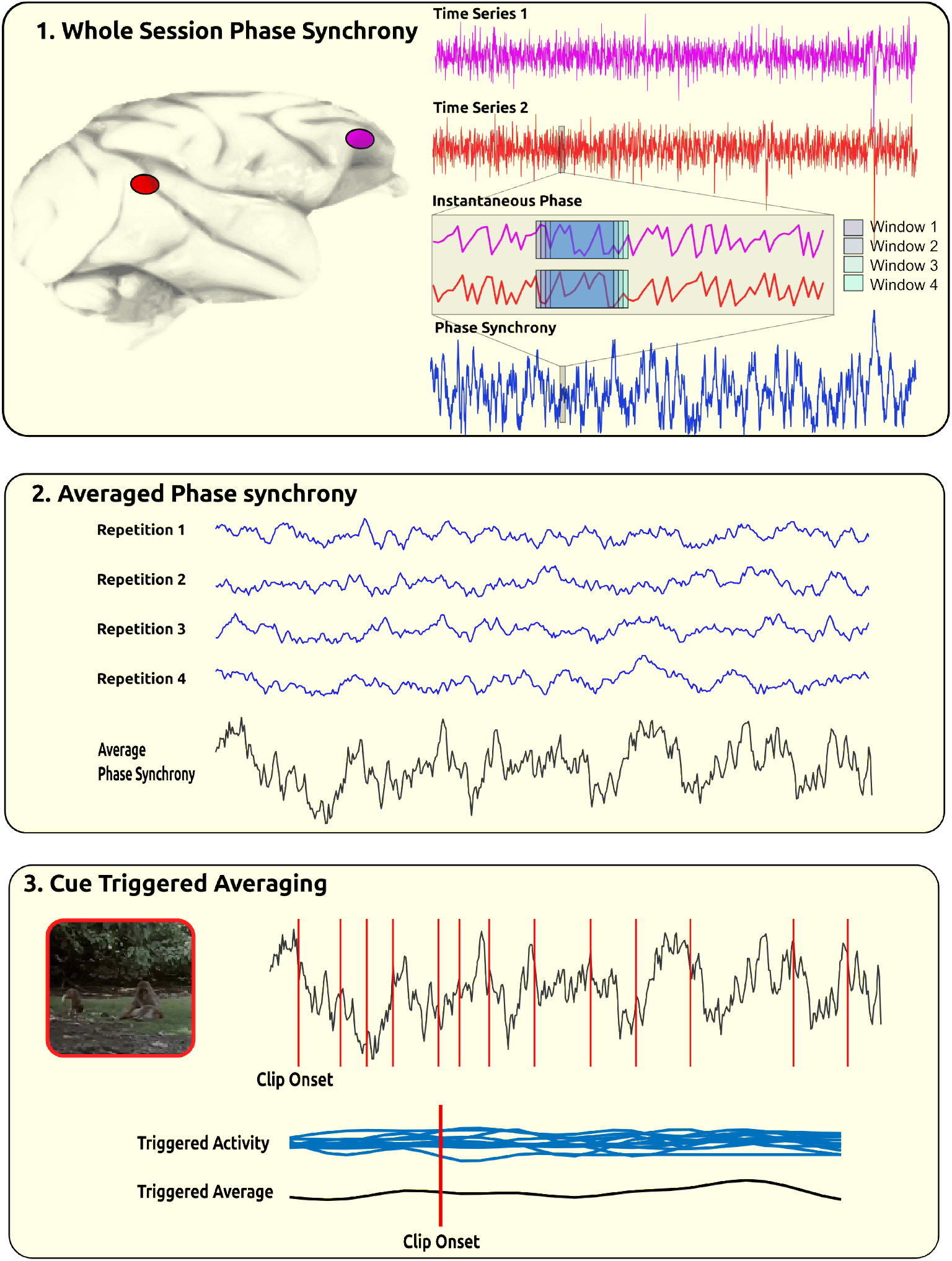
Assessing network connectivity using phase synchrony. **1.** Example whole session bold timeseries (after pre-processing & concatenation of individual runs) from one prefrontal (magenta) and one temporal (red) ROI. Phase synchrony was calculated between the instantaneous phase of both timeseries over the whole session using a window of 32 sec, overlapping the previous window by 30 sec (four overlapping windows shown inset). **2.** Whole session phase synchrony was averaged by the number of repeated runs (three or four repeats per session) in the session to yield a single time-series corresponding to the 440 volumes of unique video content. **3.** Clip onset triggered averaged were calculated from time-series aligned to the relevant clip onset (aggressive behaviour pictured) and averaged across the strongest 15% of connections between five groups (cingulate-cingulate, cingulate-temporal, premotor-cingulate, premotor-temporal and temporo-temporal).

**Figure S3.**
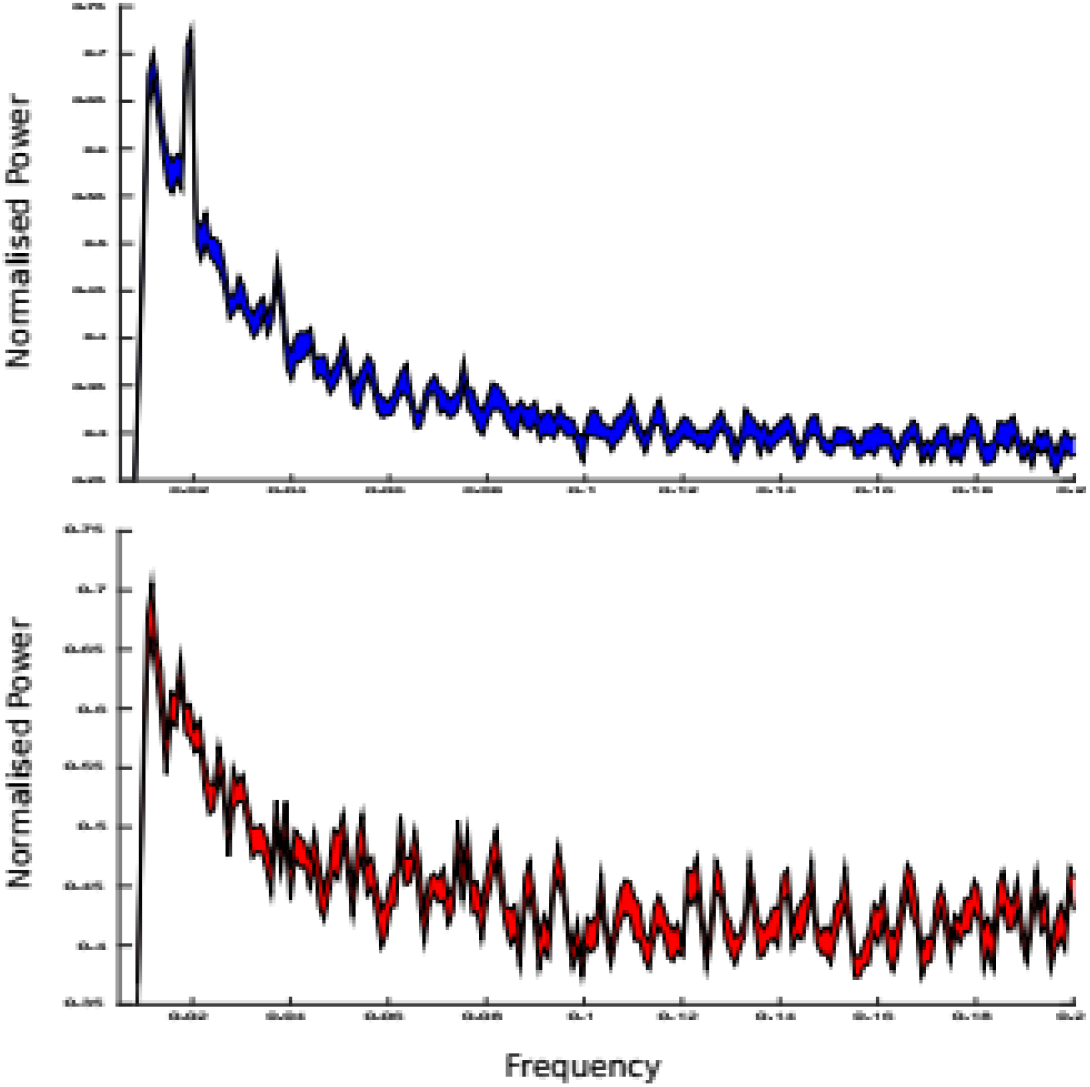
The frequency of BOLD timeseries in frontal and temporal lobes. Spectrograms calculated from the BOLD fMRI timeseries after filtering at 0.01Hz showing the average frequency content of the BOLD timeseries at ROIs in the temporal lobe (*top, blue*) and frontal lobe (*red, lower*).

**Figure S4.**
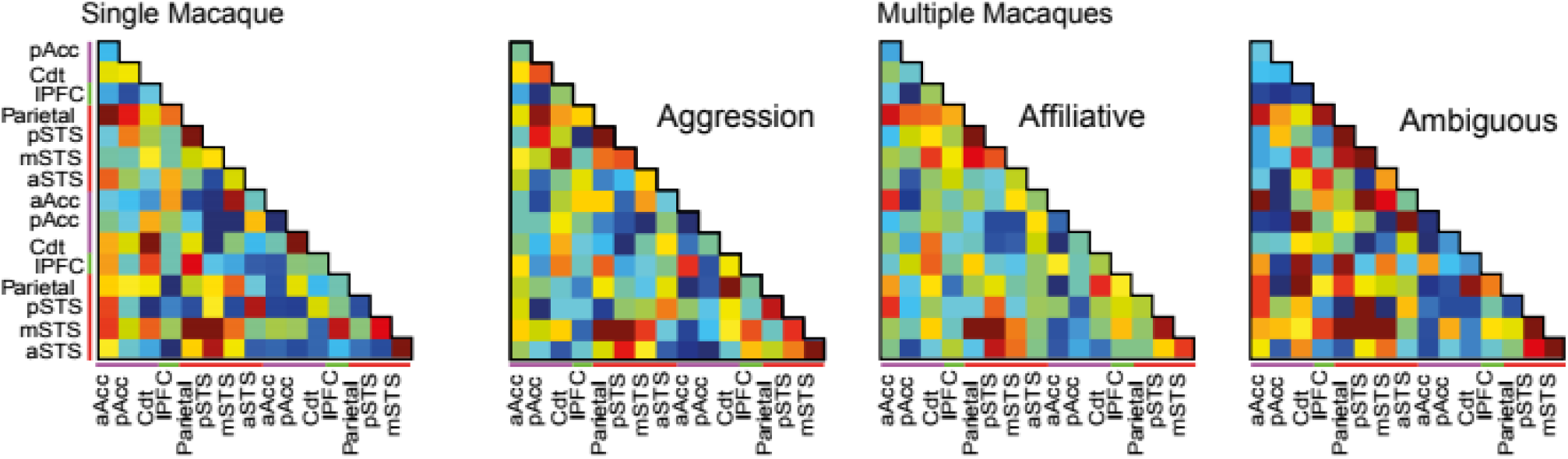
Core network connectivity associated with social video features. Average connectivity matrices calculated from scenes featuring single macaques as well as three matrices corresponding to scenes during which monkeys viewed multiple macaques engaged in three different types of behaviour (aggressive, affiliative, and ambiguous behaviour).

## REFERENCE LIST

Aharon, I., Etcoff, N., Ariely, D., Chabris, C. F., O’Connor, E., & Breiter, H. C. (2001). Beautiful faces have variable reward value: fMRI and behavioral evidence. Neuron, 32(3), 537–551. doi:10.1016/s0896-6273(01)00491-3

Ainsworth, M., Browncross, H., Mitchell, D. J., Mitchell, A. S., Passingham, R. E., Buckley, M. J., … Bell, A. H. (2018). Functional reorganisation and recovery following cortical lesions: A preliminary study in macaque monkeys. Neuropsychologia, 119, 382–391. doi:10.1016/j.neuropsychologia.2018.08.024

Alves, P. N., Foulon, C., Karolis, V., Bzdok, D., Margulies, D. S., Volle, E., & Thiebaut de Schotten, M. (2019). Subcortical Anatomy of the Default Mode Network: a functional and structural connectivity study. bioRxiv, 528679. doi:10.1101/528679

Apps, M. A., & Ramnani, N. (2014). The anterior cingulate gyrus signals the net value of others’ rewards. J Neurosci, 34(18), 6190–6200. doi:10.1523/JNEUROSCI.2701-13.2014

Arioli, M., & Canessa, N. (2019). Neural processing of social interaction: Coordinate◻based meta◻analytic evidence from human neuroimaging studies. Hum Brain Mapp. doi:10.1002/hbm.24627

Azzi, J. C., Sirigu, A., & Duhamel, J. R. (2012). Modulation of value representation by social context in the primate orbitofrontal cortex. Proc Natl Acad Sci U S A, 109(6), 2126–2131. doi:10.1073/pnas.1111715109

Baez-Mendoza, R., Harris, C. J., & Schultz, W. (2013). Activity of striatal neurons reflects social action and own reward. Proc Natl Acad Sci U S A, 110(41), 16634–16639. doi:10.1073/pnas.1211342110

Baez-Mendoza, R., van Coeverden, C. R., & Schultz, W. (2016). A neuronal reward inequity signal in primate striatum. J Neurophysiol, 115(1), 68–79. doi:10.1152/jn.00321.2015

Baumann, S., Joly, O., Rees, A., Petkov, C. I., Sun, L., Thiele, A., & Griffiths, T. D. (2015). The topography of frequency and time representation in primate auditory cortices. Elife, 4. doi:10.7554/eLife.03256

Behrens, T. E., Hunt, L. T., & Rushworth, M. F. (2009). The computation of social behavior. Science, 324(5931), 1160–1164. doi:10.1126/science.1169694

Behrens, T. E., Hunt, L. T., Woolrich, M. W., & Rushworth, M. F. (2008). Associative learning of social value. Nature, 456(7219), 245–249. doi:10.1038/nature07538

Bell, A. H., Hadj-Bouziane, F., Frihauf, J. B., Tootell, R. B., & Ungerleider, L. G. (2009). Object representations in the temporal cortex of monkeys and humans as revealed by functional magnetic resonance imaging. J Neurophysiol, 101(2), 688–700. doi:10.1152/jn.90657.2008

Bell, A. H., Malecek, N. J., Morin, E. L., Hadj-Bouziane, F., Tootell, R. B., & Ungerleider, L. G. (2011). Relationship between functional magnetic resonance imaging-identified regions and neuronal category selectivity. J Neurosci, 31(34), 12229–12240. doi:10.1523/JNEUROSCI.5865-10.2011

Bruce, V., & Young, A. (1986). Understanding face recognition. Br J Psychol, 77 (Pt 3), 305–327. Retrieved from https://www.ncbi.nlm.nih.gov/pubmed/3756376

Chang, C., Liu, Z., Chen, M. C., Liu, X., & Duyn, J. H. (2013). EEG correlates of time-varying BOLD functional connectivity. Neuroimage, 72, 227–236. doi:10.1016/j.neuroimage.2013.01.049

Chang, S. W., Gariepy, J. F., & Platt, M. L. (2013). Neuronal reference frames for social decisions in primate frontal cortex. Nat Neurosci, 16(2), 243–250. doi:10.1038/nn.3287

Chau, B. K., Sallet, J., Papageorgiou, G. K., Noonan, M. P., Bell, A. H., Walton, M. E., & Rushworth, M. F. (2015). Contrasting Roles for Orbitofrontal Cortex and Amygdala in Credit Assignment and Learning in Macaques. Neuron, 87(5), 1106–1118. doi:10.1016/j.neuron.2015.08.018

Coudé, G., Festante, F., Cilia, A., Loiacono, V., Bimbi, M., Fogassi, L., & Ferrari, P. F. (2016). Mirror Neurons of Ventral Premotor Cortex Are Modulated by Social Cues Provided by Others’ Gaze. J Neurosci, 36(11), 3145–3156; doi: 10.1523/JNEUROSCI.3220-15.2016

Cox, R. W. (1996). AFNI: software for analysis and visualization of functional magnetic resonance neuroimages. Comput Biomed Res, 29(3), 162–173. Retrieved from https://www.ncbi.nlm.nih.gov/pubmed/8812068

Dal Monte, O., Chu, C.C.J., Fagan, N.A., Chang, S.W.C. (2020). Specialized medial prefrontal-amygdala coordination in other-regarding decision preference. Nature Neuroscience. doi: 10.1038/s41593-020-0593-y

Diehl, M. M., & Romanski, L. M. (2014). Responses of prefrontal multisensory neurons to mismatching faces and vocalizations. J Neurosci, 34(34), 11233–11243. doi:10.1523/JNEUROSCI.5168-13.2014

Downing, P. E., Jiang, Y., Shuman, M., & Kanwisher, N. (2001). A cortical area selective for visual processing of the human body. Science, 293(5539), 2470–2473. doi:10.1126/science.1063414

Downing, P. E., Peelen, M. V., Wiggett, A. J., & Tew, B. D. (2006). The role of the extrastriate body area in action perception. Soc Neurosci, 1(1), 52–62. doi:10.1080/17470910600668854

Dunbar, R. I., & Shultz, S. (2007). Evolution in the social brain. Science, 317(5843), 1344–1347. doi:10.1126/science.1145463

Ebisch, S. J. H., Gallese, V., Salone, A., Martinotti, G., di Iorio, G., Mantini, D., Northoff, G. (2018). Disrupted relationship between “resting state” connectivity and task-evoked activity during social perception in schizophrenia. Schizophr Res, 193, 370–376. doi:10.1016/j.schres.2017.07.020

Fatfouta, R., Meshi, D., Merkl, A., & Heekeren, H. R. (2018). Accepting unfairness by a significant other is associated with reduced connectivity between medial prefrontal and dorsal anterior cingulate cortex. Soc Neurosci, 13(1), 61–73. doi:10.1080/17470919.2016.1252795

Ferrari, P, F., Gallese, V., Rizzolatti, G., Fogassi, L. (2003) Mirror neurons responding to the observation of ingestive and communicative mouth actions in the monkey ventral premotor cortex. Eur. J. Neurosci, 17, 1703–1714. doi:10.1046/j.1460-9528.2003.02601.x

Gallese, V., Fadiga, L., Fogassi, L., & Rizzolatti, G. (1996). Action recognition in the premotor cortex. Brain, 119(2), 593–609. doi:10.1093/brain/119.2.593

Gardner, T., Goulden, N., & Cross, E. S. (2015). Dynamic modulation of the action observation network by movement familiarity. J Neurosci, 35(4), 1561–1572. doi:10.1523/JNEUROSCI.2942-14.2015

Grabenhorst, F., Báez-Mendoza, R., Genest, W., Deco, G., & Schultz, W. (2019). Primate Amygdala Neurons Simulate Decision Processes of Social Partners. Cell, 177(4), 986–998.e915. doi:10.1016/j.cell.2019.02.042

Greve, D. N., & Fischl, B. (2009). Accurate and robust brain image alignment using boundary-based registration. Neuroimage, 48(1), 63–72. doi:10.1016/j.neuroimage.2009.06.060

Hadj-Bouziane, F., Bell, A. H., Knusten, T. A., Ungerleider, L. G., & Tootell, R. B. (2008). Perception of emotional expressions is independent of face selectivity in monkey inferior temporal cortex. Proc Natl Acad Sci U S A, 105(14), 5591–5596. doi:10.1073/pnas.0800489105

Haroush, K., & Williams, Ziv M. (2015). Neuronal Prediction of Opponent’s Behavior during Cooperative Social Interchange in Primates. Cell, 160(6), 1233–1245. doi:10.1016/j.cell.2015.01.045

Hasson, U., Nir, Y., Levy, I., Fuhrmann, G., Malach, R. (2004). Intersubject synchronization of cortical activity during natural vision. Science, 303(5664), 1634–40. doi:10.1126/science.1089506

Hill, M. R., Boorman, E. D., & Fried, I. (2016). Observational learning computations in neurons of the human anterior cingulate cortex. Nat Commun, 7, 12722. doi:10.1038/ncomms12722

Hutchison, R. M., Womelsdorf, T., Gati, J. S., Everling, S., & Menon, R. S. (2013). Resting-state networks show dynamic functional connectivity in awake humans and anesthetized macaques. Hum Brain Mapp, 34(9), 2154–2177. doi:10.1002/hbm.22058

Izuma, K., Saito, D. N., & Sadato, N. (2008). Processing of social and monetary rewards in the human striatum. Neuron, 58(2), 284–294. doi:10.1016/j.neuron.2008.03.020

Jenkinson, M., Bannister, P., Brady, M., & Smith, S. (2002). Improved optimization for the robust and accurate linear registration and motion correction of brain images. Neuroimage, 17(2), 825–841. Retrieved from https://www.ncbi.nlm.nih.gov/pubmed/12377157

Jenkinson, M., Beckmann, C. F., Behrens, T. E., Woolrich, M. W., & Smith, S. M. (2012). FSL. Neuroimage, 62(2), 782–790. doi:10.1016/j.neuroimage.2011.09.015

Jenkinson, M., & Smith, S. (2001). A global optimisation method for robust affine registration of brain images. Med Image Anal, 5(2), 143–156. Retrieved from https://www.ncbi.nlm.nih.gov/pubmed/11516708

Jimenez, A. M., Riedel, P., Lee, J., Reavis, E. A., & Green, M. F. (2019). Linking resting-state networks and social cognition in schizophrenia and bipolar disorder. Hum Brain Mapp, 40(16), 4703–4715. doi:10.1002/hbm.24731

Joly, O., Baumann, S., Balezeau, F., Thiele, A., & Griffiths, T. D. (2014). Merging functional and structural properties of the monkey auditory cortex. Front Neurosci, 8, 198. doi:10.3389/fnins.2014.00198

Jones, D. T., Vemuri, P., Murphy, M. C., Gunter, J. L., Senjem, M. L., Machulda, M. M., … Jack, C. R. Jr., (2012). Non-stationarity in the “resting brain’s” modular architecture. PLoS One, 7(6), e39731. doi:10.1371/journal.pone.0039731

Kanwisher, N., McDermott, J., & Chun, M. M. (1997). The fusiform face area: a module in human extrastriate cortex specialized for face perception. J Neurosci, 17(11), 4302–4311. Retrieved from https://www.ncbi.nlm.nih.gov/pubmed/9151747

Kolster, H., Janssens, T., Orban, G. A., & Vanduffel, W. (2014). The retinotopic organization of macaque occipitotemporal cortex anterior to V4 and caudoventral to the middle temporal (MT) cluster. J Neurosci, 34(31), 10168–10191. doi:10.1523/JNEUROSCI.3288-13.2014

Kolster, H., Mandeville, J. B., Arsenault, J. T., Ekstrom, L. B., Wald, L. L., & Vanduffel, W. (2009). Visual field map clusters in macaque extrastriate visual cortex. J Neurosci, 29(21), 7031–7039. doi:10.1523/JNEUROSCI.0518-09.2009

Koster-Hale, J., & Saxe, R. (2013). Theory of mind: a neural prediction problem. Neuron, 79(5), 836–848. doi:10.1016/j.neuron.2013.08.020

Kudo, H., & Dunbar, R. I. M. (2001). Neocortex size and social network size in primates. Animal Behaviour, 62(4), 711–722. doi:10.1006/anbe.2001.1808

Liao, W., Qiu, C., Gentili, C., Walter, M., Pan, Z., Ding, J., … Chen, H. (2010). Altered effective connectivity network of the amygdala in social anxiety disorder: a resting-state FMRI study. PLoS One, 5(12), e15238. doi:10.1371/journal.pone.0015238

Lockwood, P. L., & Wittmann, M. K. (2018). Ventral anterior cingulate cortex and social decision-making. Neurosci Biobehav Rev, 92, 187–191. doi:10.1016/j.neubiorev.2018.05.030

Mantini, D., Gerits, K., Durand, J.B., Joly, O., Simone, L., Sawamura, H., Wardak, C., Orban, G.A., Buckner, R.L., Vanduffel, W., (2011) Default mode of brain function in monkeys. J Neurosci 31(36) 12945–12962. doi:10.1523/JNEUROSCI.2318-11.2011

Mars, R. B., Neubert, F. X., Noonan, M. P., Sallet, J., Toni, I., & Rushworth, M. F. (2012). On the relationship between the “default mode network” and the “social brain”. Front Hum Neurosci, 6, 189. doi:10.3389/fnhum.2012.00189

Mars, R. B., Sallet, J., Neubert, F. X., & Rushworth, M. F. (2013). Connectivity profiles reveal the relationship between brain areas for social cognition in human and monkey temporoparietal cortex. Proc Natl Acad Sci U S A, 110(26), 10806–10811. doi:10.1073/pnas.1302956110

McCarthy, G., Puce, A., Gore, J. C., & Allison, T. (1997). Face-specific processing in the human fusiform gyrus. J Cogn Neurosci, 9(5), 605–610. doi:10.1162/jocn.1997.9.5.605

Mitchell, D. J., Bell, A. H., Buckley, M. J., Mitchell, A. S., Sallet, J., & Duncan, J. (2016). A Putative Multiple-Demand System in the Macaque Brain. J Neurosci, 36(33), 8574–8585. doi:10.1523/JNEUROSCI.0810-16.2016

Molenberghs, P., Johnson, H., Henry, J. D., & Mattingley, J. B. (2016). Understanding the minds of others: A neuroimaging meta-analysis. Neurosci Biobehav Rev, 65, 276–291. doi:10.1016/j.neubiorev.2016.03.020

Nelissen, K., Borra, E., Gerbella, M., Rozzi, S., Luppino, G., Vanduffel, W., Rizzolatti, G., Orban, G.A., (2011). Action Observation Circuits in the Macaque Monkey Cortex. J Neurosci 31(10),3743–3756. doi:10.1523/JNEUROSCI.4803-10.2011

Noonan, M. P., Sallet, J., Mars, R. B., Neubert, F. X., O’Reilly, J. X., Andersson, J. L., … Rushworth, M. F. (2014). A neural circuit covarying with social hierarchy in macaques. PLoS Biol, 12(9), e1001940. doi:10.1371/journal.pbio.1001940

Palva, J. M., Palva, S., & Kaila, K. (2005). Phase synchrony among neuronal oscillations in the human cortex. J Neurosci, 25(15), 3962–3972. doi:10.1523/JNEUROSCI.4250-04.2005

Peelen, M. V., Wiggett, A. J., & Downing, P. E. (2006). Patterns of fMRI activity dissociate overlapping functional brain areas that respond to biological motion. Neuron, 49(6), 815–822. doi:10.1016/j.neuron.2006.02.004

Peirce, J. W. (2007). PsychoPy--Psychophysics software in Python. J Neurosci Methods, 162(1-2), 8–13. doi:10.1016/j.jneumeth.2006.11.017

Pinsk, M. A., Arcaro, M., Weiner, K. S., Kalkus, J. F., Inati, S. J., Gross, C. G., & Kastner, S. (2009). Neural representations of faces and body parts in macaque and human cortex: a comparative FMRI study. J Neurophysiol, 101(5), 2581–2600. doi:10.1152/jn.91198.2008

Platt, M. L., Seyfarth, R. M., & Cheney, D. L. (2016). Adaptations for social cognition in the primate brain. Philos Trans R Soc Lond B Biol Sci, 371(1687), 20150096. doi:10.1098/rstb.2015.0096

Popivanov, I. D., Jastorff, J., Vanduffel, W., & Vogels, R. (2012). Stimulus representations in body-selective regions of the macaque cortex assessed with event-related fMRI. Neuroimage, 63(2), 723–741. doi:10.1016/j.neuroimage.2012.07.013

Power, J. D., Barnes, K. A., Snyder, A. Z., Schlaggar, B. L., & Petersen, S. E. (2012). Spurious but systematic correlations in functional connectivity MRI networks arise from subject motion. Neuroimage, 59(3), 2142–2154. doi:10.1016/j.neuroimage.2011.10.018

Rabany, L., Diefenbach, G. J., Bragdon, L. B., Pittman, B. P., Zertuche, L., Tolin, D. F., … Assaf, M. (2017). Resting-State Functional Connectivity in Generalized Anxiety Disorder and Social Anxiety Disorder: Evidence for a Dimensional Approach. Brain Connect, 7(5), 289–298. doi:10.1089/brain.2017.0497

Rizzolatti, G., & Sinigaglia, C. (2010). The functional role of the parieto-frontal mirror circuit: interpretations and misinterpretations. Nat Rev Neurosci, 11(4), 264–274. doi:10.1038/nrn2805

Romanski, L. M., & Diehl, M. M. (2011). Neurons responsive to face-view in the primate ventrolateral prefrontal cortex. Neuroscience, 189, 223–235. doi:10.1016/j.neuroscience.2011.05.014

Rosenblum, M. G., Pikovsky, A. S., & Kurths, J. (1996). Phase synchronization of chaotic oscillators. Phys Rev Lett, 76(11), 1804–1807. doi:10.1103/PhysRevLett.76.1804

Rubinov, M., & Sporns, O. (2010). Complex network measures of brain connectivity: uses and interpretations. Neuroimage, 52(3), 1059–1069. doi:10.1016/j.neuroimage.2009.10.003

Rudebeck, P. H., Buckley, M. J., Walton, M. E., & Rushworth, M. F. (2006). A role for the macaque anterior cingulate gyrus in social valuation. Science, 313(5791), 1310–1312. doi:10.1126/science.1128197

Russ B.E., Leopold D.A. (2015). Functional MRI mapping of dynamic visual features during natural viewing in the macaque. Neuroimage, 109, 84–94. doi: 10.1016/j.neuroimage.2015.01.012

Sallet, J., Mars, R. B., Noonan, M. P., Andersson, J. L., O’Reilly, J. X., Jbabdi, S., …. Rushworth, M. F. (2011). Social network size affects neural circuits in macaques. Science, 334(6056), 697–700. doi:10.1126/science.1210027

Sallet, J., Mars, R. B., Noonan, M. P., Neubert, F. X., Jbabdi, S., O’Reilly, J. X., … Rushworth, M. F. (2013). The organization of dorsal frontal cortex in humans and macaques. J Neurosci, 33(30), 12255–12274. doi:10.1523/JNEUROSCI.5108-12.2013

Sapey-Triomphe, L.-A., Centelles, L., Roth, M., Fonlupt, P., Hénaff, M.-A., Schmitz, C., & Assaiante, C. (2017). Deciphering human motion to discriminate social interactions: a developmental neuroimaging study. Social Cognitive and Affective Neuroscience, 12(2), 340–351. doi:10.1093/scan/nsw117

Saxe, R., & Kanwisher, N. (2003). People thinking about thinking people. The role of the temporo-parietal junction in “theory of mind”. Neuroimage, 19(4), 1835–1842. Retrieved from https://www.ncbi.nlm.nih.gov/pubmed/12948738

Saxe, R., Xiao, D. K., Kovacs, G., Perrett, D. I., & Kanwisher, N. (2004). A region of right posterior superior temporal sulcus responds to observed intentional actions. Neuropsychologia, 42(11), 1435–1446. doi:10.1016/j.neuropsychologia.2004.04.015

Scalaidhe, S. P., Wilson, F. A., & Goldman-Rakic, P. S. (1999). Face-selective neurons during passive viewing and working memory performance of rhesus monkeys: evidence for intrinsic specialization of neuronal coding. Cereb Cortex, 9(5), 459–475. doi:10.1093/cercor/9.5.459

Schulke, O., Bhagavatula, J., Vigilant, L., & Ostner, J. (2010). Social bonds enhance reproductive success in male macaques. Curr Biol, 20(24), 2207–2210. doi:10.1016/j.cub.2010.10.058

Seidlitz, J., Sponheim, C., Glen, D., Ye, F. Q., Saleem, K. S., Leopold, D. A., … Messinger, A. (2018). A population MRI brain template and analysis tools for the macaque. Neuroimage, 170, 121–131. doi:10.1016/j.neuroimage.2017.04.063

Sergent, J., Ohta, S., & MacDonald, B. (1992). Functional neuroanatomy of face and object processing. A positron emission tomography study. Brain, 115 Pt 1, 15–36. doi:10.1093/brain/115.1.15

Sescousse, G., Li, Y., & Dreher, J. C. (2015). A common currency for the computation of motivational values in the human striatum. Social Cognitive and Affective Neuroscience, 10(4), 467–473. doi:10.1093/scan/nsu074

Sliwa, J., & Freiwald, W. A. (2017). A dedicated network for social interaction processing in the primate brain. Science, 356(6339), 745–749. doi:10.1126/science.aam6383

Tsao, D. Y., Moeller, S., & Freiwald, W. A. (2008). Comparing face patch systems in macaques and humans. Proc Natl Acad Sci U S A, 105(49), 19514–19519. doi:10.1073/pnas.0809662105

Tsao, D. Y., Schweers, N., Moeller, S., & Freiwald, W. A. (2008). Patches of face-selective cortex in the macaque frontal lobe. Nat Neurosci, 11(8), 877–879. doi:10.1038/nn.2158

Van Essen, D. C., Glasser, M. F., Dierker, D. L., & Harwell, J. (2012). Cortical parcellations of the macaque monkey analyzed on surface-based atlases. Cereb Cortex, 22(10), 2227–2240. doi:10.1093/cercor/bhr290

Viviano, J. D., Buchanan, R. W., Calarco, N., Gold, J. M., Foussias, G., Bhagwat, N., … Social Processes Initiative in Neurobiology of the Schizophrenia, G. (2018). Resting-State Connectivity Biomarkers of Cognitive Performance and Social Function in Individuals With Schizophrenia Spectrum Disorder and Healthy Control Subjects. Biol Psychiatry, 84(9), 665–674. doi:10.1016/j.biopsych.2018.03.013

Wagner, D. D., Haxby, J. V., & Heatherton, T. F. (2012). The representation of self and person knowledge in the medial prefrontal cortex. Wiley Interdiscip Rev Cogn Sci, 3(4), 451–470. doi:10.1002/wcs.1183

Wagner, D. D., Kelley, W. M., Haxby, J. V., & Heatherton, T. F. (2016). The Dorsal Medial Prefrontal Cortex Responds Preferentially to Social Interactions during Natural Viewing. Journal of Neuroscience, 36(26), 6917–6925. doi:10.1523/jneurosci.4220-15.2016

Watson, Karli K., & Platt, Michael L. (2012). Social Signals in Primate Orbitofrontal Cortex. Current Biology, 22(23), 2268–2273. doi:10.1016/j.cub.2012.10.016

Wittmann, M. K., Kolling, N., Faber, N. S., Scholl, J., Nelissen, N., & Rushworth, M. F. (2016). Self-Other Mergence in the Frontal Cortex during Cooperation and Competition. Neuron, 91(2), 482–493. doi:10.1016/j.neuron.2016.06.022

Wittmann, M. K., Lockwood, P. L., & Rushworth, M. F. S. (2018). Neural Mechanisms of Social Cognition in Primates. Annu Rev Neurosci, 41, 99–118. doi:10.1146/annurev-neuro-080317-061450

Woolrich, M. W., Behrens, T. E., Beckmann, C. F., Jenkinson, M., & Smith, S. M. (2004). Multilevel linear modelling for FMRI group analysis using Bayesian inference. Neuroimage, 21(4), 1732–1747. doi:10.1016/j.neuroimage.2003.12.023

Yoshida, K., Saito, N., Iriki, A., & Isoda, M. (2011). Representation of Others’ Action by Neurons in Monkey Medial Frontal Cortex. Current Biology, 21(3), 249–253. doi:10.1016/j.cub.2011.01.004

Yoshida, K., Saito, N., Iriki, A., & Isoda, M. (2012). Social error monitoring in macaque frontal cortex. Nat Neurosci, 15(9), 1307–1312. doi:10.1038/nn.3180

Zhang, Y., Brady, M., & Smith, S. (2001). Segmentation of brain MR images through a hidden Markov random field model and the expectation-maximization algorithm. IEEE Trans Med Imaging, 20(1), 45–57. doi:10.1109/42.906424

Zhu, H., Qiu, C., Meng, Y., Yuan, M., Zhang, Y., Ren, Z., … Zhang, W. (2017). Altered Topological Properties of Brain Networks in Social Anxiety Disorder: A Resting-state Functional MRI Study. Scientific reports, 7, 43089. doi:10.1038/srep43089

